# miR160 controls class III peroxidase via *StARF10/17* and couples salicylic acid and ROS signaling networks in potato hypersensitive response to potato virus Y

**DOI:** 10.64898/2026.03.04.709499

**Authors:** Maja Križnik, Tjaša Lukan, Maja Zagorščak, Anna Coll Rius, Karmen Pogačar, Nastja Marondini, Katja Stare, Valentina Levak, Jitka Široká, Ondřej Novák, Kristina Gruden

## Abstract

- Hypersensitive response (HR) is an effector-triggered immune response leading to pathogen restriction and local cell death. Small RNAs (sRNAs), mediating RNA silencing, have a well-established role in antiviral immunity, however, their role in HR has not been addressed to date.
- We applied a spatially resolved sRNAomics approach to uncover changes at the sRNA level and the dynamics of sRNA–gene regulatory networks in potato HR response to potato virus Y (PVY).
- We show that miR160 is repressed in cells adjacent to HR lesions in the resistant cultivar, but not in PVY-sensitive, salicylic acid (SA)-depleted plants. Beyond its canonical regulation of *StARF10* and *StARF17*, miR160 controls a broader regulatory network spanning auxin signaling, cell-wall remodeling, redox homeostasis, and tuberization. Elevated miR160 levels promote susceptibility-associated traits, reduce thermotolerance, and suppress genes strongly activated in HR.
- Mechanistically, we identify a novel miR160–*StARF10/StARF17*–*StPRX28* module that links miR160 repression to the induction of a class III apoplastic peroxidase during HR, with SA acting upstream. Together, these findings establish miR160 as a crucial integrator of SA–auxin–ROS signaling pathways that tunes development, defense, and heat resilience, and highlight miR160 as a promising target for fine-tuning tolerance to multiple stresses with minimal yield loss.

## Introduction

In nature, plants are constantly challenged by a plethora of microbial pathogens and pests. During an arms race, plants have evolved complex fine-tuned defense mechanisms. Pattern-triggered immunity (PTI) relies on the recognition of pathogen-/damage-/microbe-/herbivore-associated molecular patterns (PAMPs/DAMPs/MAMPs/HAMPs) by plant membrane-located pattern recognition receptors (PRRs), comprising from receptor-like kinases (RLKs) and receptor-like proteins (RLPs) (Ngou et al., 2022a). However, some adapted pathogens are able to circumvent PTI by production of virulence proteins, i.e. effectors. Plants, in turn, have evolved additional layers of defense that depend on the recognition of pathogen effectors by intracellular nucleotide-binding/leucine rich-repeat receptors (NLRs), which are often encoded by *R* genes. Activation of NLR immune receptors results in effector-triggered immunity (ETI) (Ngou et al., 2022b). The ETI response commonly culminates in programmed cell death (PCD) at infection sites, referred to as the hypersensitive response (HR) (Künstler *et al*., 2016). Increasing number of studies have illustrated that PTI and ETI are interdependent and mutually potentiate each other to achieve robust resistance against pathogens (Yu *et al*., 2024).

In addition to protein-coding genes acting as key regulators, increasing evidence highlights the importance of small non-coding RNAs in regulating plant immunity (Qiao *et al*., 2021; Lopez-Gomollon & Baulcombe, 2022). Plants produce two major classes of small RNAs (sRNAs), microRNAs (miRNAs) and small interfering RNAs (siRNAs), which are distinguished by their modes of origin and biogenesis (Axtell, 2013). sRNAs are generally 20–24 nucleotides in length and mediate post-transcriptional gene silencing by guiding mRNA cleavage, translation inhibition; or transcriptional gene silencing through DNA modification in a sequence-specific manner (Qiao *et al*., 2021). Of the sRNAs, miRNAs, have become recognized as integral components of the functional regulatory networks orchestrating plant growth, development and response to biotic and abiotic stresses by mainly regulating the expression level and dynamics of transcription factors during the life cycle of plants (Song *et al*., 2019; Shen *et al*., 2024). Accumulating evidence highlights important roles for miRNAs as fine-tuning regulators of diverse aspects of PTI and ETI (Deng *et al*., 2018; Song *et al*., 2021; Feng *et al*., 2021). Numerous miRNAs have been implicated in modulating the delicate balance between plant growth and immunity by regulating the biosynthesis and signaling pathways of plant hormones, particularly auxin and gibberellin, which exhibit antagonistic interactions with salicylic acid (SA)-mediated immunity against pathogens.

For example, in Arabidopsis the induction of miR393 by the flagellin-derived peptide flg22 contributes to PTI by silencing multiple auxin receptors, including *TIR1*, *AFB2*, and *AFB3*, consequently restricting the growth of virulent *Pseudomonas syringae* pv. tomato (Pto) DC3000 (Navarro *et al*., 2006). We also showed that in the tolerant response, the increased expression of sRNAs negatively regulating gibberellic acid (GA) biosynthesis and signaling coincided with increased expression of immune receptor genes in response to potato virus Y (PVY) in potato (Križnik *et al*., 2017). Similarly, the upregulation of miR160, targeting auxin response factors, has been associated with enhanced immunity against the blast fungus *Magnaporthe oryzae* in rice (Li *et al*., 2014).

The potato (*Solanum tuberosum* L.) is a crucial global staple crop, renowned for its significant economic importance. Among the many viruses infecting potato, PVY (genus *Potyvirus*) is the most economically important and devastating virus infecting cultivated potato, causing potato tuber necrotic ringspot disease (Chikh-Ali *et al*., 2020). This disease poses a significant threat to potato production, with yield reductions of up to 70% and a decline in tuber quality, rendering the affected crops unsuitable for market sale (Karasev & Gray, 2013; Chikh-Ali *et al*., 2020). In potato, there are two main types of resistance to PVY: extreme resistance, conferred by the *Ry* genes, and HR, conferred by the *Ny* genes (reviewed in Kogovšek & Ravnikar, 2013). The potato cultivar Rywal carries the *Ny-1* gene and develops HR, leading to formation of necrotic lesions on inoculated leaves and arrest of virus spread (Szajko *et al*., 2008; Baebler *et al*., 2014). Similarly to some other pathosystems, in Rywal HR-associated programmed cell death is uncoupled from resistance (Lukan *et al*., 2018). HR is a highly spatiotemporally regulated process, characterized by viral spread being confined to only a limited number of cells adjacent to the infection site, and then successfully arrested by the host’s defense machinery (Lukan *et al*., 2018).

Although some insights into spatial regulation of transcription in the HR to PVY have already been gained (Lukan *et al*., 2020), post-transcriptional regulation mediated by sRNAs at such a fine spatial scale has not yet been elucidated. Addressing this knowledge gap, we adapted an approach involving the analysis of small, spatially resolved tissue sections, separately capturing PVY-induced response in cells immediately adjacent to the cell death zones (i.e. lesions), as well as the surrounding regions at different times after inoculation. This enabled fine-scale profiling of sRNA-mediated regulation during the HR and revealed that miR160 is a vital molecular hub in the SA–auxin–reactive oxygen species (ROS) network balancing developmental and stress pathways.

Functionally, miR160 was found to be a positive regulator of growth and development in potato. However, in HR response to PVY, miR160 was repressed 3 days post infection (dpi) and 5 dpi. Similarly, reduced miR160 levels were also associated with enhanced tolerance to heat stress. Collectively, these findings indicate that miR160 repression shifts the regulatory balance from growth toward defense, thereby promoting effective stress responses both against viral pathogens and heat.

## Materials and Methods

### Generation of miR160 transgenic lines

All miR160 transgenic plants used in this work were generated in the Rywal genetic background. For generation of potato miR160 inducible overexpression plants (*MIR160a-OE4*, *MIR160a-OE18*), the miR160a precursor sequence was amplified from genomic DNA (gDNA) of potato cv. Rywal with primers harboring *XhoI* or *SpeI* restriction enzyme recognition sites (pre-miR160a_OE_F: GTCAC**CTCGAG**GATCATGTAGAGATTATGAATTTAAAGAGC and pre-miR160a _OE_R: GTCGG**ACTAGT**CCATAACCTAGTTTAGATCTCAACA. The amplicon was purified using the Wizard® SV Gel and PCR Clean-Up System and ligated into the pTA7002 vector (Aoyama & Chua, 1997), under the control of a glucocorticoid-inducible promoter. The resulting construct was transformed into *Escherichia coli* One Shot™ TOP10 (Invitrogen) and verified by colony PCR using KAPA Taq PCR Kit (Kapa Biosystems) according to manufacturer’s protocol. Plasmids were isolated from positive colonies using the GeneElute Plasmid Miniprep Kit (Sigma) and subjected to Sanger sequencing (Eurofins Genomics) (Table S1). Primers used for colony PCR and sequencing were as follows: pTA7002_F (ACCTCGATCGAGATCTTCGCA) and pTA7002_R (GTGTGGGCAATGAAACTGATGC). Transgenic miR160 knockdown lines *mir160a* and *mir160b* were generated via CRISPR technology. The construct used for generation of *mir160b* is described in (Lukan *et al*., 2022). Verified constructs transformed into *Agrobacterium tumefaciens* strain LBA4404 were used to generate miR160 overexpression and knockdown lines by stable transformation as described before (Lukan *et al*., 2023). miR160 knockdown lines were genotyped to determine types of mutations and exclude off-target effect (Fig. S1, Table S1) as explained in Lukan et al. (2022). The overexpression lines were grown and sub-cultured on regeneration medium with hygromycin selection (final concentration 20 mg/l) and knockdown lines on medium with kanamycin selection (final concentration 50 mg/l) in a growth chamber (PSI; Photon Systems Instruments) under controlled environmental conditions 21/19 °C (day/night) with a long-day (16 h) photoperiod of light (light intensity 90 µmol m^-2^ s^-1^) and 40% relative humidity.

### Growth conditions

Non-transgenic potato cv. Rywal (NT), miR160 overexpression (OE) and miR160 knockdown lines, SA-depleted *NahG* transgenic line, all in Rywal genetic background, were used in this study. Plants were propagated from stem node tissue cultures and transferred to soil two weeks after node segmentation, where they were kept in PSI growth chambers under controlled environmental conditions at 21/19 °C (day/night) with a long-day (16 h) photoperiod of light (light intensity 120 µmol m^-2^ s^-1^) and 55% relative humidity. *Nicotiana benthamiana* plants were grown from seeds and kept in growth chambers under the same conditions with light intensity 80 µmol m^-2^ s^-1^.

### Plant treatments

After three weeks of growth in soil, plants were inoculated with PVY^N-Wilga^ (PVY^N–Wi^; accession no. EF558545) or with PVY^N605^, tagged with green fluorescent protein (PVY-N605(123)-GFP) (Lukan *et al*., 2023) or mock-inoculated as described in Baebler et al. (2009). To induce miR160 overexpression, miR160 OE lines were treated with dexamethasone (Dex) foliar spray solution containing 30 μM Dex and 0.01 % (v/v) Tween-20, or control spray solution without Dex (control), 3 h prior to virus/mock inoculation, then 3 h after treatment and subsequently every 24 h until sampling. To account for potential Dex effects, all NT, *NahG* and miR160 knockdown plants used in the same experiments as OE lines were also Dex-treated until sampling. *N. benthamiana* plants were inoculated with PVY-N605(123)-GFP or mock-inoculated after three weeks of growth in soil. For heat-stress experiments, plants were grown in soil for three or five weeks and then were treated with Dex every 24 h. One day after the first Dex application, plants assigned to the heat-stress condition were transferred to a separate PSI growth chamber at 30/28 °C (day/night), whereas control plants were maintained at 22/19 °C (day/night), as described in the “Growth conditions” section.

### Plant phenotypisation

Heat-treated and control potato plants grown in soil were phenotyped after 10 days of heat treatment using the PlantScreen™ SC system (PSI). The photosynthetic parameters were assessed using kinetic chlorophyll fluorescence imaging, following a measurement protocol aligned with previously established methods (Abdelhakim *et al*., 2024). Hyperspectral imaging was used to capture spectral reflectance patterns of the plants, providing indicators of physiological status. Reflected radiation was recorded across a broad wavelength range and converted into hyperspectral image data. Processed images were displayed using false-color scales, and relevant spectral parameters were subsequently extracted. All raw images were processed automatically, and trait values were extracted using the PlantScreen™ Analyzer software (PSI). For guard cell size analysis, leaf discs were collected using biopunch (Miltex® Biopsy Punch with Plunger, ID 4.0 mm, OD 4.36 mm) and immediately fixed in 3% glutaraldehyde prepared in 0.1 M phosphate buffer (pH 6.8) and subsequently used for confocal imaging. To assess the effect of miR160 on tuberization, tubers from normally grown miR160 overexpression and knockdown plants were weighed after 8 weeks and 2.5 months of growth in soil. Differences between the lines were evaluated using Dunn’s test for multiple comparisons, with *P* values adjusted using Holm’s method to control the family-wise error rate. PVY-infected and mock-inoculated *N. benthamiana* plants were analyzed for disease symptoms and PVY distribution throughout the plants using the Newton 7.0 BIO FT500 imaging system (Vilber) after two weeks post inoculation. GFP emission from the GFP-labeled PVY clone was monitored in the 480 nm excitation channel and F-550 emission filter. Images were acquired using EvolutionCapt edge software.

### sRNA and RNA Sequencing

For sRNA-Seq and RNA-Seq, lesions were sampled from PVY^N-Wi^-inoculated leaves at 3 dpi and 5 dpi. From each time point, twenty tissue sections encompassing the lesion and its immediately adjacent cells (section A), and twenty surrounding tissue sections (section B), measuring approximately 0.5 × 0.5 mm at 3 dpi, and 1 × 1 mm at 5 dpi, were collected and pooled separately (pool A and pool B). For mock-inoculated plants, small sections were prepared by cutting the narrow bands of the tissue between veins from distal to proximal site of the leaf to take the possible positional effect of the lesions into account. Sections of mock-inoculated plants were also pooled separately (pool M). Three pools were collected per each group. The pools were stored in 100 µl of RNAlater RNA Stabilization Solution (Thermo Fisher Scientific). Total RNA was extracted with TRIzol (Invitrogen) and Direct-zol RNA MicroPrep Kit, DNAse treated and purified with RNA Clean & Concentrator kit (both Zymo Research) according to the manufacturer’s instructions with prior homogenization using Tissue Lyser (Qiagen). The RNAs quantity and purity were assessed using a Bioanalyser 2100 and an RNA 6000 Pico LabChip Kit (both Agilent). Small RNA and mRNA libraries were prepared and sequenced at LC Sciences and Novogene at HiSeq4000 and NovaSeq6000 platforms (both Illumina) using 50-bp single-end for sRNA-Seq and 150-bp paired-end reads for RNA-Seq. sRNA-Seq analysis was performed as described previously (Križnik *et al*., 2021). Briefly, reads were preprocessed to remove adapter sequences and low-quality reads (Phred quality score < 20) using cutadapt (https://cutadapt.readthedocs.io/en/stable/index.html) and FASTX-Toolkit (http://hannonlab.cshl.edu/fastx_toolkit/index.html). Resulting reads were mapped with no mismatches to annotated potato miRNAs from miRbase (release 22) (Kozomara *et al*., 2019), and to previously identified potato small RNAs (novel miRNAs, miRNA variants and phasiRNAs (Križnik *et al*., 2017)). Detected sRNAs were then quantified and subjected to differential expression analysis using limma-voom approach as previously described (Križnik *et al*., 2021). For RNA-Seq, reads were quantified with Salmon using a transcriptome built from the *S. tuberosum* Phureja DM1-3 v6.1 genome (Pham *et al*., 2020) and the unified potato annotation (Zagorščak *et al*., 2024). Differential expression analysis was conducted using edgeR (v3.42.4) and limma (v3.56.2) (Ritchie *et al*., 2015; Chen *et al*., 2025). Salmon-derived counts were first scaled by the transcript-specific overdispersion estimates (function Overdispersion), then TMM-normalized and transformed using the voom function (Law *et al*., 2014), followed by empirical Bayes moderation. For both sRNAs and genes, features with Benjamini–Hochberg FDR-adjusted *P*-values < 0.05 were considered significantly differentially expressed. Gene Set Enrichment Analysis (GSEA) was performed on TMM-normalized count values using GSEA v4.3.3 (Subramanian *et al*., 2005). Gene sets with FDR-adjusted Q-values < 0.05 were considered significantly enriched in up- or down-regulated genes and visualized using adapted gseaFromStats function from the biokit v0.1.1 package (Steenwyk *et al*., 2022).

### qPCR

For spatial miRNA and gene expression analysis by RT-qPCR, individual tissue sections A, B and M were collected at 5 dpi and stored as described above for RNA-Seq. To measure PVY accumulation in upper non-inoculated leaves, 50 mg of leaf tissue were collected at 18 dpi. To analyze miRNA and gene expression under normal growth conditions (i.e. non-infected plants), 50 mg of leaves were collected after three weeks of growth in soil followed by five days of Dex treatment. All sampled leaves were snap-frozen in liquid nitrogen. Tissue sections and leaves were homogenized using Tissue Lyser (Qiagen), followed by RNA isolation using TRIzol and Direct-zol RNA MicroPrep Kit (for small tissue sections) and Direct-zol RNA MiniPrep Plus Kit (for leaf tissue). Stem-loop RT-qPCR was used to quantify the expression of miRNAs. DNAse treated RNA was reverse transcribed using SuperScript III First-Strand Synthesis System following the manufacturer’s protocol with the following modifications: 100 U SuperScript III enzyme per reaction, 0.5 × stem-loop primer concentration (both Thermo Fisher Scientific), and use of a pulsed reverse transcription protocol (Varkonyi-Gasic and Hellens 2011; Križnik et al. 2017). For individual sections, RT product (5 µl) was pre-amplified using TaqMan PreAmp Master Mix (2x) and PreAmp miRNA Primer Mix (5x) (both Thermo Fisher Scientific) in a 25-µl PCR reaction according to the manufacturer’s instructions. All qPCR reactions were performed in 5 µl volumes in duplicates and two dilutions (10- and 50-fold) per sample using 2.75 µl TaqMan Universal Master Mix II and 0.25 µl TaqMan MicroRNA Assays (both Applied Biosystems). miR160 expression was quantified and normalized to the expression of endogenous control stu-miR167a-5p.1 as described previously (Križnik *et al*., 2017) using quantGenius (http://quantgenius.nib.si; (Baebler *et al*., 2017)). For gene expression analyses, RNA was DNAse treated (RNase-Free DNase Set, Qiagen) and reverse transcribed using High-Capacity cDNA Reverse Transcription Kit (Applied Biosystems) according to manufacturer’s instructions. All qPCR reactions were performed in 5 µl volumes in duplicates and two dilutions (10- and 50-fold) per sample using FastStart Universal Probe Master Mix (Roche). Gene expression was quantified by normalization to the two validated reference genes *StCox* and *StEf-1* with quantGenius (Baebler *et al*., 2017). Viral quantity was analyzed in tissue sections sampled at 5 dpi, and in leaf tissue of the upper non-inoculated leaves sampled at 18 dpi. Sample preparation and qPCR analysis were performed as described above using PVY as a target gene. For primer and probe information, see Table S2. To quantify the magnitude of group differences in relative miRNA and gene expression, Hedges’ g was calculated with bias correction for small sample sizes (effsize v0.8.1; Torchiano, 2020). Uniformity, dispersion, and outliers were assessed using diagnostic simulations implemented in the DHARMa v0.4.7 package (Hartig, 2024). Subsequently, spatial data—either log_2_-transformed or 97th-percentile-scaled—were analyzed by ANOVA followed by Tukey’s Honest Significant Difference (HSD) test to identify group differences, whereas non-spatial data were log₂-transformed prior to two-tailed Welch’s t-test analysis. For visualization, log_2_-transformed values were scaled relative to the corresponding control group mean (scaled value = log_2_(measurement) − mean(log_2_(control)), yielding the log_2_ fold change (log_2_FC) values that facilitate intuitive interpretation of up- or down-regulation across experimental conditions. Figures were generated using ggplot2 v4.0.0 (Wickham, 2016) and ggpubr v0.6.1.1 (Kassambara, 2025).

### Hormonal measurements

For hormonal content measurements 100 sections (2 mm x 2 mm) encompassing both section A and B, and sections from mock-inoculated leaves (M), were sampled from inoculated leaves at 5 dpi. Three pools (∼12 mg each) were collected per each group. Tissue was snap-frozen in liquid nitrogen. Hormones were extracted and analyzed by liquid chromatography coupled with tandem mass spectrometry as described before (Široká *et al*., 2022). Briefly, the samples were simultaneously homogenized and extracted in 1 mL of 1 mol L^-1^ formic acid in 10% aqueous methanol with the addition of internal standards ([2H6]-JA, [2H2]-(-)-JA-Ile, [2H5]-OPDA, [13C6]-IAA, [2H6]-ABA (OlChemIm, Czech Republic), and [2H4]-SA (Sigma Aldrich, USA) using 5 ceria-stabilized zirconium oxide beads in MM 400 mixer mill (Retsch GmbH, Haan, Germany) (29 Hz, 10 min, precooled holders). Subsequently, the samples were centrifuged (16 000 g, 10 min, 8 °C) and 200 μL of the supernatant was purified on reverse phase microSPE columns assembled from C18 and SDB-XC sorbents. The analyses were performed on a 1290 Infinity liquid chromatography system coupled to a 6490 Triple Quadrupole mass spectrometer (Agilent Technologies, USA) and the data were processed in MassHunter Quantitative B.09.00 software (Agilent Technologies, USA). To account for values below the limit of detection (LOD), measurements were imputed as one-tenth of the minimum observed value, hormone-wise. Differences in hormonal levels were assessed using Welch’s ANOVA followed by Games–Howell post-hoc test (implemented via rstatix v0.7.2) on log₁₀-transformed data. Effect sizes were estimated using Wilcoxon’s r statistics. Results were visualised using ggplot2 v4.0.0 and ggpubr v0.6.1. 1.

### Construction of sRNA-target regulatory networks

For construction of sRNA-target regulatory network, potato sRNAs (miRNAs, siRNAs) identified previously (miRbase, release 22; Križnik *et al*., 2017) were linked with their mRNA targets through *in silico* predictions and degradome-Seq data. *In silico* identification of sRNA-target interactions was performed using the psRNATarget v2 (Dai *et al*., 2018) against the unified potato transcriptome (Zagorščak *et al*., 2024), following previously proposed stringent parameters (Zhang *et al*., 2013). For degradome-Seq data analysis, four potato degradome libraries of PVY-infected and mock-inoculated samples (GEO accession no. GSE84966) generated previously (Križnik *et al*., 2017) were re-analyzed using CleaveLand4 (Addo-Quaye *et al*., 2009) with the unified potato transcriptome (Zagorščak *et al*., 2024). Parameters were as described in the original study (Križnik *et al*., 2017). sRNA–target interactions were integrated with expression data to identify active sRNA–target regulatory modules in HR to PVY at 3 dpi. Network was constructed in Cytoscape 3.10.2 (Shannon *et al*., 2003) (Dataset S1). Differentially expressed sRNAs in HR to PVY at 3 dpi were additionally analyzed for functional overrepresentation in biological pathways using the MapMan software (Usadel *et al*., 2009) using MapMan4 ontology (Schwacke *et al*., 2019). Functional bin assignments were assigned based on sRNA target annotations predicted through Mercator4 v6 via the web platform (https://www.plabipd.de/mercator_main.html). Additionally, to identify miR160-regulated genes downstream of ARFs, miR160-target regulatory networks were constructed based on genes showing significant differential expression in miR160 overexpression and knockdown plants compared with NT plants under normal and PVY-infected conditions. Network was constructed in Cytoscape 3.10.2 (Shannon *et al*., 2003) (Dataset S2).

### Transactivation assay

To validate predicted transcriptional regulation *in planta*, we performed transient transactivation assay according to Lasierra & Prat (2018). For this, the full length *StARF10* and *StARF17* cDNA were amplified from cv. Rywal and cloned by pENTR™ Directional TOPO® Cloning Kit (Invitrogen) into pENTR D-TOPO cloning vector and recombined into the β-estradiol inducible vector containing GFP (pABinGFP; (Bleckmann *et al*., 2010)) using the Gateway LR Clonase II Enzyme Kit (Thermo Fisher Scientific) to obtain the *pABinGFP_StARF10* and *pABinGFP_StARF17* constructs. Promoter sequence of *StPRX28* (∼1 kb) was amplified from cv. Rywal and inserted into the pENTR D-TOPO vector (Invitrogen) and subsequently recombined through LR reaction into the pGWB435 Gateway vector (Nakagawa *et al*., 2007) as described previously (Tomaž *et al*., 2023), inserting the promoter upstream of a luciferase reporter (*FLuc)*. All primer pairs used in the cloning procedure are listed in Table S3. Leaf discs of *N. benthamiana* plants transiently co-expressing the promoter *FLuc* reporter and β-estradiol inducible transcription factor effector cassettes, in combination with RNA silencing suppressor p19, were sampled 4 days post agroinfiltration. For effector construct activation, β-estradiol was added to a final concentration of 500 nM. No inducer was added to half of the leaf discs, used as controls for basal activity. For each effector construct, three independent experiments were performed. Luminescence activity was recorded in 10 min intervals with Centro LB963 Luminometer (Berthold Technologies). Pointwise group comparisons were performed using pairwise Wilcoxon tests (implemented via rstatix v0.7.2; (Kassambara, 2023)) and visualised using ggplot2 v4.0.0. package (Wickham, 2016).

### Confocal Imaging

To follow virus spread around the cell death zones, PVY-N605(123)-GFP inoculated leaves were analyzed using confocal microscope (Leica TCS LSI macroscope with Plan APO 5x objective or Leica Stellaris 8 with HC PL FLUOTAR 10x objective (both Leica Microsystems, Germany)). GFP was excited with the 488 nm laser and the emission was followed in the window between 505 and 530 nm. The background chlorophyll (Chl) fluorescence was excited with the 488 nm laser and the emission was measured in the window between 690 and 750 nm. The fluorescence was monitored on the adaxial side of leaf discs containing the lesions, sampled with biopunch (Miltex® Biopsy Punch with Plunger, ID 4.0 mm, OD 4.36 mm). Fluorescence emissions were collected sequentially through two channels, GFP and Chl fluorescence. Regions of interest were scanned unidirectionally with frame average 2 and scan speed 400 Hz. Z-stack size was adjusted between 6 to 12 steps. The images were processed using Leica Las X software (Leica Microsystems) to obtain maximum projections from z-stacks for both channels. For transactivation assays, the transcription factor production was confirmed by measuring fluorescent tag with the Leica TCS LSI macroscope with Plan APO 20× objective, using the settings described above. For observation of tissue organization in the leaves, Leica Stellaris 8 with HC PL FLUOTAR 10x objective was used. Adaxial leaf tissue was visualized via Chl autofluorescence. Regions of interest were scanned unidirectionally with frame average 2 and scan speed 600 Hz or bidirectionally with line average 2 and scan speed 400 Hz. Z-stack size was adjusted to cover the palisade tissue. For guard cell size analysis, Leica TCS LSI macroscope equipped with an ACS APO 20×/0.60 IMM objective was used. The abaxial leaf surface was examined using a Leica DFC7000 T digital camera for brightfield imaging, where the region of interest was either scanned by acquiring a z-stack (10 steps, with step sizes ranging from 1.5 to 7.0 µm) or by capturing an image. Images were processed using Leica Las X software to obtain maximum projections from z-stacks. Guard cell size was quantified by measuring the length and width of each guard cell pair using the software’s measurement tool. Guard cell size was calculated using an ellipse-based formula: π×(length/2) ×(width/2), as previously described (Zhang *et al*., 2023). Differences in guard cell size among normally grown potato lines were evaluated using Dunn’s test for multiple comparisons, with *P* values adjusted using Holm’s method to control the family-wise error rate. For the laser power, PIN opening and detector gain please check metadata available on Zenodo (10.5281/zenodo.18797111).

### *In silico* sequence and structural analysis

The amino acid sequences of Arabidopsis, tomato (*Solanum lycopersicum* L.) and potato defined as peroxidases in PLAZA 5.0 (Van Bel *et al*., 2022) were used to construct a phylogenetic tree. To determine the optimal number of sequence clusters, affinity propagation clustering (v1.4.13; (Bodenhofer *et al*., 2011)) was conducted on mutual pairwise similarities obtained through protein length scaled Levenshtein distances (v0.9.15; (Van Der Loo, 2014)). Circular dendrogram was constructed using the R (v4.4.1) package dendextend (v1.19.0; (Galili, 2015)). Protein domain prediction was performed with ExPASy Prosite (https://prosite.expasy.org/) and InterPro (https://www.ebi.ac.uk/interpro/). SignalP v6.0 was used for signal peptide predictions (Teufel *et al*., 2022). DeepLoc v2.0 (Thumuluri *et al*., 2022) and CELLO v2.5 (Yu *et al*., 2006) were used for cellular localization prediction. Predictions of transcription factor binding motifs in promoter sequences of *MIR160* loci and promoters of miR160-regulated genes were performed with PlantPAN 4.0. (Chow *et al*., 2024). Alignment of pre-miR160a and pre-miR160b sequences derived from the *MIR160a* and *MIR160b* loci was performed using MAFFT v7 (Katoh *et al*., 2019). Predicted secondary structures of pre-miR160a and pre-miR160b were generated using RNAfold implemented in the ViennaRNA Package v2.1.5 (Lorenz *et al*., 2011). Structural models of StARF10 and StARF17 DNA binding domain (∼350 aa; (Cancé *et al*., 2022)) were generated with AlphaFold3 (Abramson *et al*., 2024). The top-ranked models for each ARF were selected. The ChimeraX molecular visualization program was used for visual analysis and calculation of Coulombic electrostatic potential (ESP).

## Results

### miR160 is downregulated in the cells adjacent to the cell death zone of hypersensitive response to potato virus Y

In this study, we focused on unravelling the dynamics of sRNA regulatory networks in ETI of potato to PVY. Plants of cv. Rywal exhibit typical HR resistance to PVY, effectively restricting the virus within the inoculated leaves while exhibiting the characteristic development of necrotic lesions 3–5 days post infection (dpi) (Fig. 1a). Given the spatial intricacies inherent in the HR, where the virus is known to be confined near to the infection site but escapes the cell death zone (Lukan *et al*., 2018), we employed spatially resolved approach to decipher the role of sRNAs in the process. Tissue samples from the lesions and immediately adjacent cells (section A) and from the surrounding regions (section B) were collected at the early visible (3 dpi) and fully developed (5 dpi) lesion stages using our recently developed sampling protocol for spatial precision transcriptomics (Lukan *et al*., 2020). In addition, protocols for sRNA extraction and quantification were adapted to accommodate minute quantities, enabling both targeted and non-targeted spatial analyses.

**Fig. 1:**
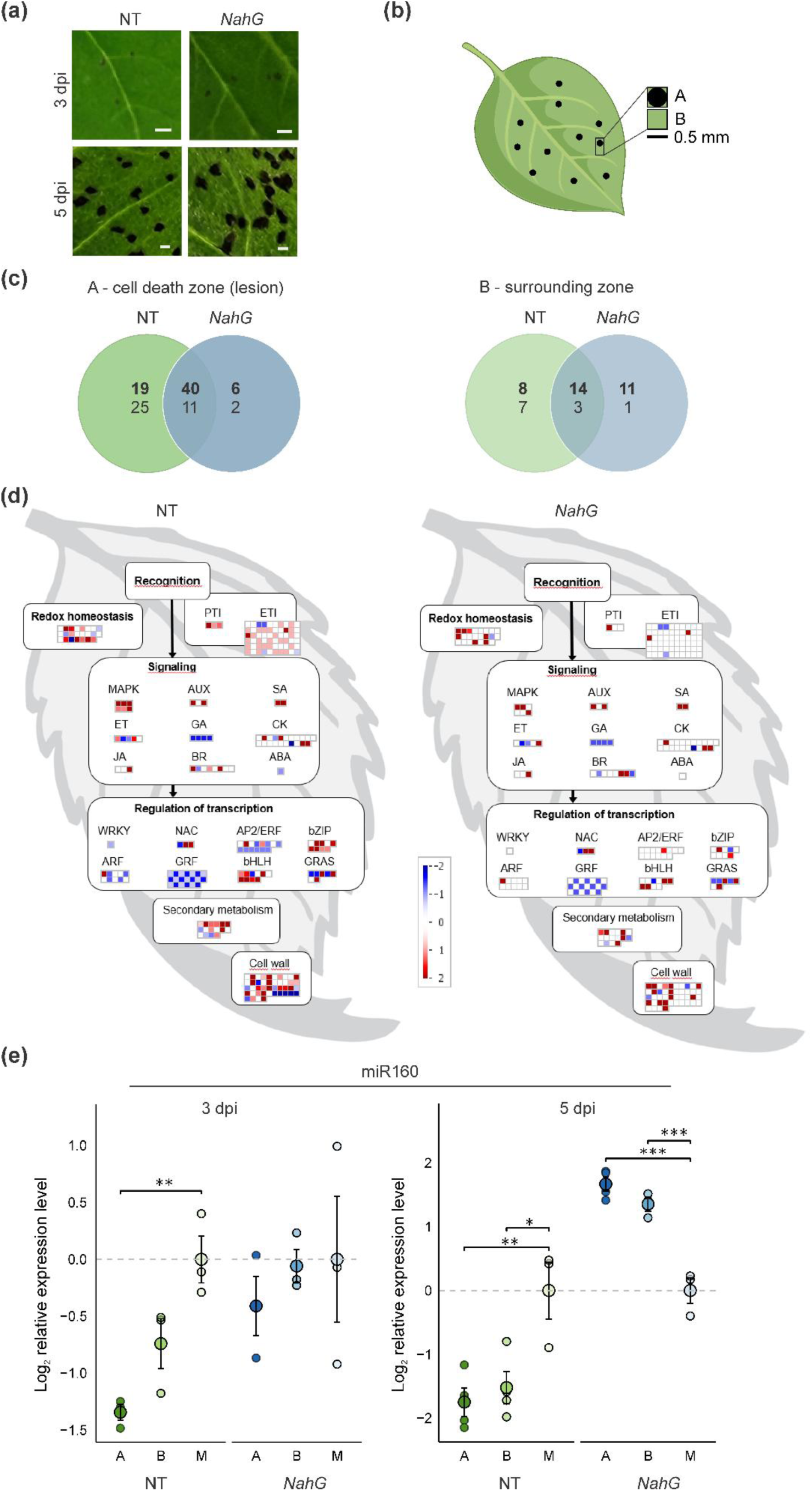
Spatial regulation of sRNAs in HR to PVY in potato. (a) PVY-induced lesions on inoculated leaves of resistant cv. Rywal (NT) and sensitive SA-depleted *NahG* plants at early (3 days post infection (dpi)) and fully developed stages (5 dpi). Photographs were taken using a LED light panel to enhance lesion visibility. Scale bar: 1 mm. (b) Sampling scheme at 3 dpi: tissue sections including lesion and its immediately adjacent cells (section A) and surrounding region (section B). (c) Numbers of unique and common differentially expressed sRNAs (miRNAs and phasiRNAs) in section A and section B at 3 dpi in response to PVY in NT and *NahG* plants. Bold format numbers indicate upregulated sRNAs, while regular format numbers denote downregulated sRNAs. (d) sRNAs target multiple immune- and developmental-related components in resistant NT. Visualization of differentially expressed sRNAs identified in section A at 3 dpi in NT and *NahG* plants, according to the function of their predicted targets. Each square represents one target gene and colors represent log_2_ ratios of sRNA expression between PVY- and mock-inoculated plants (red - upregulated; blue - downregulated; FDR-adjusted *P* < 0.05). MapMan4 ontology Bins: Redox homeostasis (10), Pathogen.pattern-triggered immunity (PTI) machinery (26.10.1), Pathogen.effector-triggered immunity (ETI) machinery (26.10.2), MAPK (27.14), AUX (11.2), SA (11.8), ET (11.5), GA (11.6), CK (11.4), JA (11.7), BR (11.3), ABA (11.1), WRKY (15.5.7.5), NAC (15.5.7.1), AP/ERF (15.5.7.6), bZIP (15.5.1.1), ARF (15.5.9.1), GRF (15.5.2.6), bHLH (15.5.1.2), GRAS (15.5.10.2), secondary metabolism (9), cell wall (21). (e) miR160 shows differential spatial regulation in section A between NT and *NahG* plants. Relative miR160 expression level was determined in sections A and B at 3 dpi and 5 dpi, at the stage of early and fully developed lesions, respectively, together with mock-inoculated tissue sections (M). Relative miR160 levels were log_2_-transformed and for visualization purpose scaled to the arithmetic mean of the corresponding control (M). Biological replicates (small circles) and arithmetic mean (large, dark filled circles) ± standard error of the mean are shown. Asterisks denote statistically significant differences based on Tukey’s HSD test (\**P*<0.05, \*\**P*<0.01, *** *P*< 0.001).

First, we explored the sRNA responses in the early visible lesions that develop 3 dpi following inoculation with PVY (Fig. 1a,b). To identify sRNAs associated with efficient viral arrest, we compared the sRNÁs response of non-transgenic PVY-resistant plants of cv. Rywal (hereafter referred to as NT) with response of sensitive SA-depleted *NahG* transgenic plants. *NahG* plants exhibit impaired resistance to PVY, characterized by uncontrolled systemic viral spread and formation of systemic lesions (Fig. S2). We found that in HR, PVY infection triggers a localized increase in sRNAs cells immediately adjacent to the cell death zone (section A), which diminishes in the surrounding tissue (section B) (Fig. 1c, Fig. S3, Table S4). Construction of sRNA-target regulatory network identified modules with differential spatial activation between NT and *NahG* (Fig. 1d; Table S5; Dataset S1). We found that sRNAs downregulated in cells adjacent to cell death zones of resistant NT plants target transcripts encoding developmental transcription factors, such as *ARFs*, and *AP2/ERFs* (Fig. 1d, Table S5). Among sRNAs examined, stu-miR160a,b-5p (hereafter referred to as miR160) was significantly downregulated at 3 dpi in cells immediately adjacent to the cell death zone of resistant NT but not in *NahG* plants (Fig. 1e, Table S5). To further investigate the spatial dynamics of the miR160 response, we analyzed its expression at 5 dpi, a time point when lesions were fully developed and the virus was effectively contained in NT but spread in *NahG* plants (Fig. S4). We found that miR160 levels were further reduced in NT in cells adjacent to the cell death zone and in surrounding tissue relative to mock (Fig. 1e, Table S6), while *NahG* plants exhibited marked upregulation of miR160 in these regions. This upregulation correlated with impaired resistance phenotype and larger lesion size of *NahG* plants compared to NT (Fig. 1a, Table S6), therefore we hypothesized that miR160 may be involved in SA-deficiency–driven susceptibility to PVY.

### Unveiling the miR160 downstream signaling network in potato

miR160 is an evolutionarily conserved miRNA, encoded by two *MIR* genes in potato, *MIR160a* and *MIR160b* (miRBase, Križnik *et al*., 2017), and processed from the 5′ arms of pre-miR160a and pre-miR160b precursor transcripts (Fig. S5). We have previously shown that miR160 directs the cleavage of two *ARF* genes in potato, *StARF10* and *StARF17* (Križnik *et al*., 2017). To investigate broader miR160 function through direct or indirect regulation, we generated miR160 overexpression (OE) potato lines in cv. Rywal genetic background (Fig. 2a). To avoid potential developmental abnormalities or lethality associated with constitutive miRNA expression in potato, we generated plants using an inducible glucocorticoid-based system (Aoyama & Chua, 1997) where expression of pre-miR160a precursor is induced by the external application of dexamethasone (Dex). *MIR160a-OE4 and MIR160a-OE18* lines were selected for their highest accumulation of miR160 (Fig. 2a,b). To additionally examine the effect of miR160 depletion, we generated CRISPR-Cas9 knockdown mutants of *MIR160a* and *MIR160b* genes (*mir160a*, *mir160b*; Fig. 2a, Fig. S1, Table S1; Lukan *et al*., 2022). In *mir160a* mutants, miR160 levels were reduced to ∼20% of native values, whereas in *mir160b* mutants they were reduced to ∼50% (Fig. 2b, Table S7). Paralogous *MIR160a* gene appears to be more potent generator of miR160 than *MIR160b*, as its silencing leads to stronger reduction of miR160 levels (Fig. 2b, Table S7). No viable *mir160a mir160b* double mutants were recovered, suggesting that the combined loss of both genes is lethal.

**Fig. 2.**
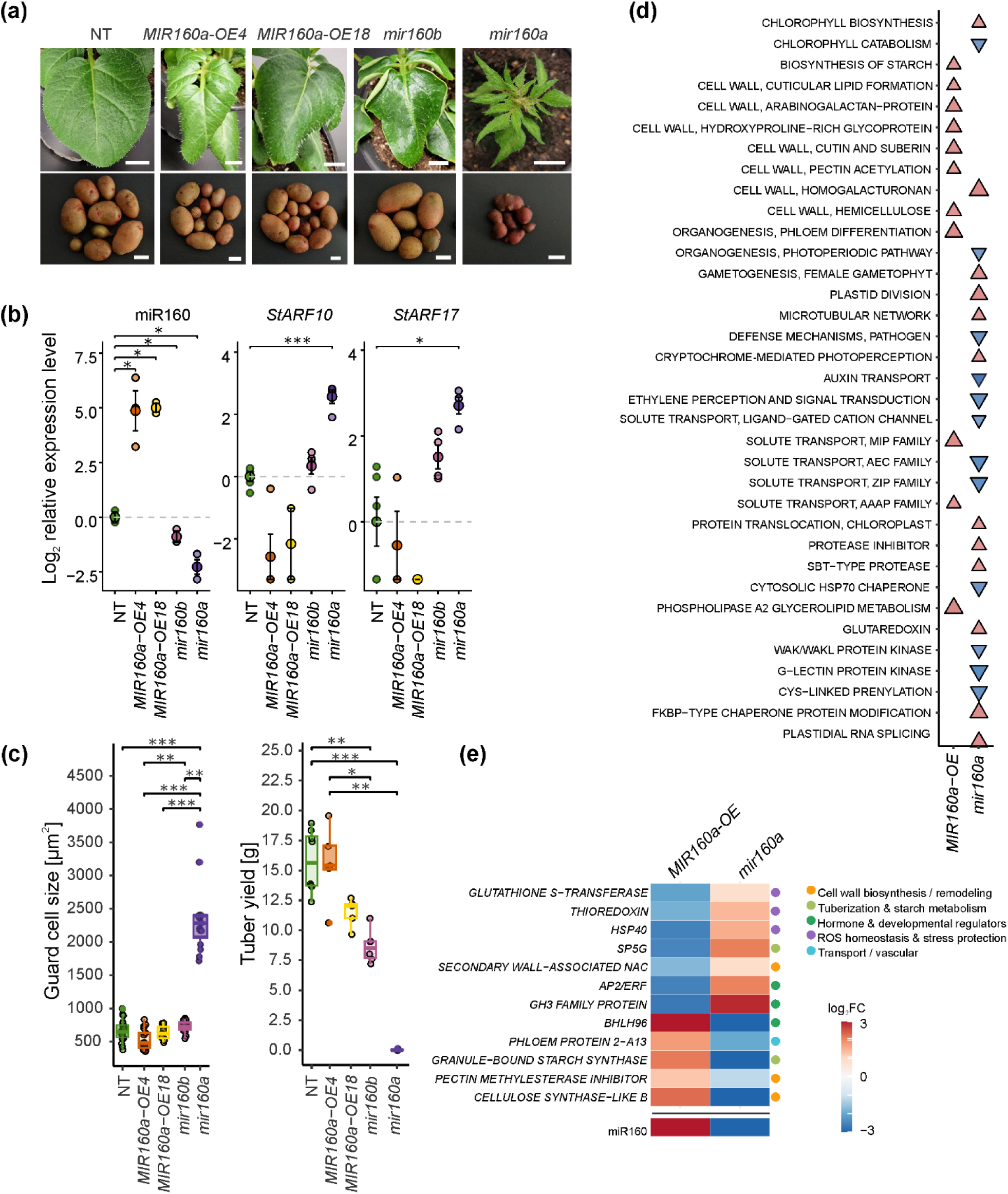
Changes in miR160 levels impact redox homeostasis and cell wall metabolism, in addition to auxin signaling. (a) Phenotypic differences of miR160 overexpression (*MIR160a-OE4, MIR160a-OE18*) and miR160 knockdown lines (*mir160a*, *mir160b*) compared with non-transgenic (NT) plants. Plants were grown in soil for three weeks and treated with dexamethasone (Dex) for 5 days prior to leaf imaging. Tubers were imaged after 10 weeks of growth. Scale bar: 1 cm. (b) Relative expression levels of miR160 and their targets *StARF10* and *StARF17*. Relative expression levels were log_2_ transformed and Welch’s t-test was applied to evaluate differences between groups. For visualization purposes values were further scaled to the arithmetic mean of the reference group (NT). Biological replicates (small circles) and arithmetic mean (large, dark filled circles) ± standard error of the mean are shown. Asterisks denote statistically significant differences (\**P*<0.05, \*\**P*<0.01, *** *P*< 0.001). (c) Guard cell size and tuber yield of miR160 transgenic lines compared to NT. Asterisks denote statistically significant differences (\**P*<0.05, \*\**P*<0.01, *** *P*< 0.001) determined using Dunn’s method for multiple comparisons with Holm’s *P*-value adjustment. (d) Impact of miR160 on regulation of multiple biological processes. MapMan4 ontology BINs were used to generate gene sets. Only statistically significant gene sets enriched in comparison with NT (FDR Q-value < 0.05) determined by gene set enrichment analysis (GSEA) are presented. ”red triangles“- induced processes, ”blue triangles“- repressed processes, blanks denote that a current process was not statistically significantly enriched. For the full list see Table S11. (e) Subset of newly identified miR160-regulated genes showing inverse differential expression between overexpression and knockdown plants (red - upregulated; blue - downregulated; FDR-adjusted *P* < 0.05). For the full list of miR160-regulated genes and their descriptions, see Table S10.

*MIR160a-OE* lines showed no significant differences in growth rate compared to NT, as expected due to the inducible overexpression system employed. However, when the plants were treated for induction of miR160 overexpression downward leaf curling of leaf margins (i.e. epinasty) was observed after 5 days of treatment (Fig. 2a). Conversely, the *mir160b* mutants exhibited upward leaf curling (i.e. hyponasty) and a wavy/undulating leaf surface (Fig. 2a). Strong reduction of miR160 accumulation in *mir160a* knockdown plants lead to substantial growth retardation and impaired lamina development, resulting in small sickle-shaped leaves (Fig. 2a). At the cellular level, mesophyll cells of *mir160a* mutants exhibited irregular shapes and orientations, indicative of uneven, anisotropic growth (Fig. S6). In addition, *miR160a* plants developed larger guard cells than NT (Fig. 2c, Fig. S7, Table S8). Strongly diminished miR160 levels in *mir160a* plants also impaired below-ground development, with root and tuber growth significantly reduced relative to NT. *mir160b* plants, however, showed only a slight reduction in tuber yield (Fig. 2c, Fig. S8, Table S9). Reflecting the extent of phenotypic changes observed, non-targeted RNA-Seq approach revealed differential regulation of more than 8,000 genes in *mir160a* mutants, whereas 828 genes were differentially expressed in miR160 OE plants compared to NT (Table S10). The lower number of differentially expressed genes in OE plants likely reflects that miR160 overexpression was induced for only five days in fully grown plants, whereas the knockdown mutants were depleted of miR160 from the early stages of development.

We further explored the effects of miR160 modulation at the transcriptome level. In miR160-deficient plants, genes associated with auxin transport, ethylene signaling, and chlorophyll catabolism were significantly downregulated (Fig. 2d, Table S11). Conversely, in miR160 OE plants, processes linked to solute transport, organogenesis, and cell wall organization—each tightly intertwined with auxin signaling—were upregulated (Fig. 2d, Table S11). Consistent with the positive role of miR160 in growth and development, besides *StARF10* and *StARF17* miR160 also represses the auxin-responsive *StGH3* (encoding an auxin-conjugating enzyme) and a tuberization repressor *StSP5G*. Moreover, miR160 downregulates genes involved in ROS homeostasis, including *glutathione S-transferase (GSTU)*, a *thioredoxin family gene,* and a *Hsp40* heat stress–responsive chaperone (Fig. 2e, Table S10). All of these miR160-regulated genes except *GSTU,* contain ARF-binding motifs in their promoters, positioning them directly downstream of the ARF regulatory network (Fig. 2e, Table S10, Table S12a, Dataset S2).

### miR160 does not break *R*-gene resistance but it represses several genes that are induced in hypersensitive response

To study whether elevated miR160 compromises resistance in HR, miR160 overexpression plants were inoculated with GFP-tagged PVY and examined for viral spatial distribution in infected leaves near the entry site using confocal microscopy at 6 dpi, which corresponds to the stage when the virus spread is already successfully blocked in NT, but not in sensitive *NahG* (Fig. S4). miR160 overexpression did not influence virus spread; PVY remained confined close to the infection site, similar to NT (Fig. S10). Evidence that elevated levels of miR160 do not compromise resistance was further supported by a three-week monitoring period, during which no PVY accumulation in upper non-inoculated leaves was detected (Table S13).

To investigate further whether miR160 is involved in regulating defense signaling processes downstream of *Ny-1* gene activation, we analyzed the responses of miR160 OE lines and NT at the fully developed lesion stage (5 dpi) in section A and section B (Fig. 3a). We identified additional miR160-regulated genes (Fig. 3b, Table S14, Dataset S2). Most of miR160-repressed genes were strongly upregulated in A and B sections of NT plants. This set includes genes implicated in cellular energy metabolism, redox homeostasis, and tuberization (Fig. 3b, Table S14). In addition, we quantified levels of *StHSP70*, previously proposed to be modulated by miR160 in Arabidopsis (Lin *et al*., 2018), in individual tissue sections. Indeed, *StHSP70* was significantly downregulated in miR160 OE plants compared with NT (Fig. 3c, Table S15). We also quantified the expression of *peroxidase 28* (*StPRX28*), which was strongly induced in section A of NT plants (Fig. 3c, Table S15). Similarly to *StHSP70*—and with a larger effect—*StPRX28* expression was significantly reduced in miR160 OE lines at the lesion margins and in the surrounding tissues compared with NT (Fig. 3c, Fig. S11, Table S15). Together, our findings reveal that *StPRX28*, a PVY-induced and spatially regulated peroxidase, is a novel regulatory target of miR160, whose induction coincides with spatially graded downregulation of miR160 in lesions of NT (Fig. 1e), which may function to relieve repression of *StPRX28* in HR to PVY.

**Fig. 3.**
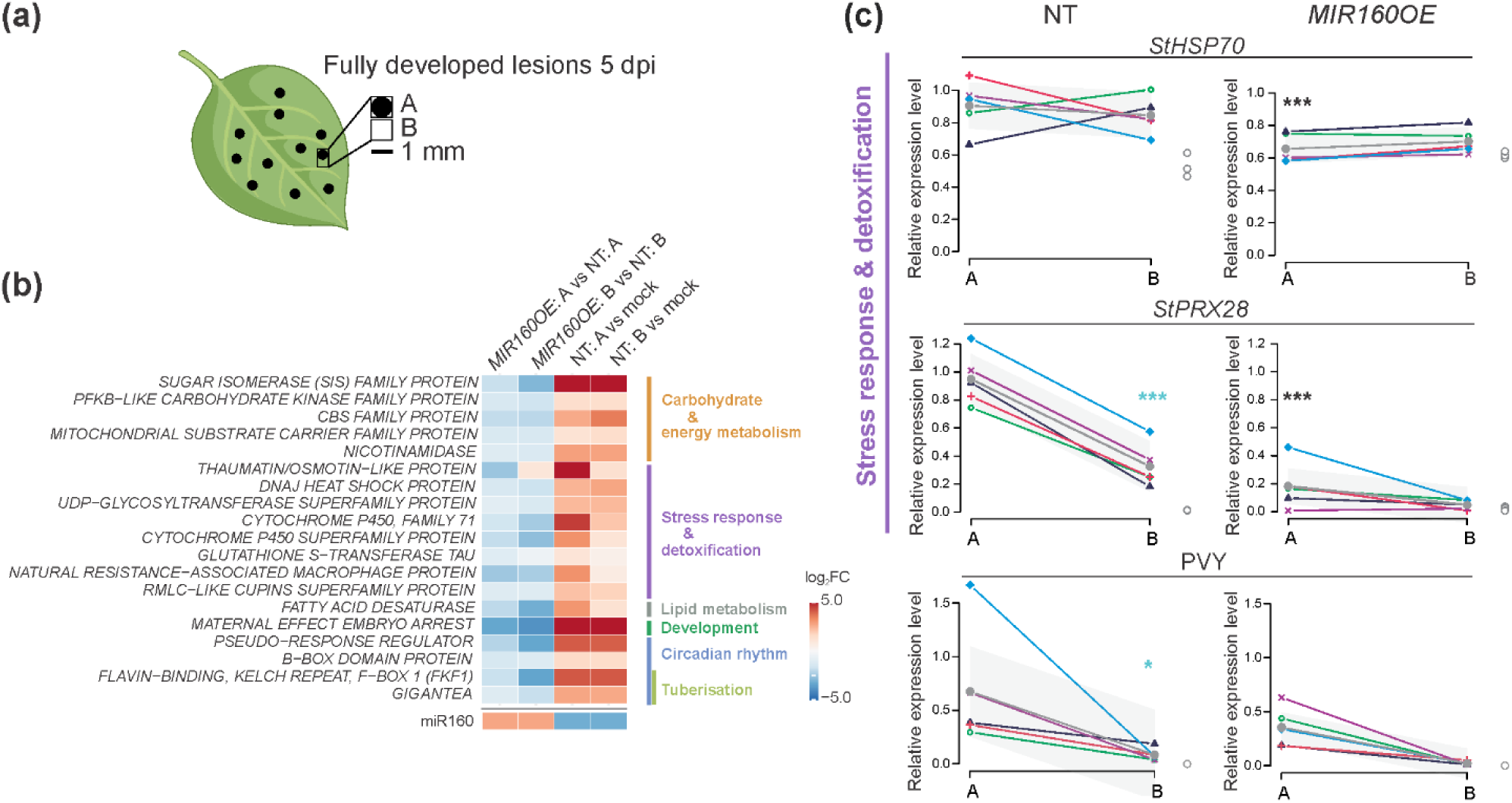
miR160 suppresses several genes upregulated in HR-conferred resistance. (a) Sampling scheme at 5 days post infection (dpi): tissue sections including cells immediately adjacent to the cell death zones (section A) and surrounding regions (section B). (b) Transcriptional profiles of the genes negatively regulated by miR160 and upregulated in NT (non-transgenic) following inoculation with PVY at the stage of fully developed lesions (5 dpi) in A and B sections. The values represent log_2_FC; red-upregulated, blue-downregulated. The last row of the heatmap shows the spatial response of miR160 under the same set of comparisons. For the full list of miR160-regulated genes and their descriptions, see Table S10. (c) Spatial response of *StHSP70*, *StPRX28* and PVY levels in individual A and B tissue sections. Expression values in mock (M)-inoculated tissue sections are shown as empty circles at the end of x-axis. *MIR160OE* plots correspond to the *MIR160a-OE4* line. The results for independent transgenic line *MIR160a-OE18* are similar to *MIR160a-OE4* line and are given in Fig. S11 and Table S15. Spatial profile models are shown as thick black lines, with 95% confidence interval bands presented in grey. Relative gene expression values of individual lesions are presented with colored symbols connected by a line. Black asterisks indicate statistically significant differences in gene expression in miR160 OE plants in sections A and B relative to NT, whereas blue asterisks indicate statistically significant differences between sections A and B within each genotype. Differences between and within genotypes were assessed by Tukey HSD test (\**P*<0.05, \*\**P*<0.01, *** *P*< 0.001).

### miR160 negatively regulates Class III apoplastic peroxidase *StPRX28* via StARF10 and StARF17

To elucidate the link between miR160 and *StPRX28*, we first performed *in silico* miRNA target prediction analyses and degradome-Seq analyses to test for direct interaction between miR160 and *StPRX28*. However, neither approach identified sufficient complementarity to support miR160-guided cleavage or translational repression, making direct interaction unlikely. To explore other options, we analyzed *cis*-regulatory elements in the *StPRX28* promoter region (2 kb upstream of the start codon) of the potato genome (*S. tuberosum* Phureja DMv6). The *StPRX28* promoter shows enrichment of zinc finger homeodomain (ZF-HD) binding motifs and harbors binding sites for WRKY, and bZIP/TGA transcription factors implicated in SA signaling, together with AP2/ERF binding motifs associated with the ethylene signaling pathway (Fig. S12a, Table S12b). Notably, we identified two ARF-binding motifs (AuxRE, TGTCTC) located at 567 bp (+ strand) and at 600 bp (- strand) upstream of the transcription start site (Fig. S12b, Table S12b). AlphaFold-3 models of StARF10 and StARF17 DNA binding domains (DBD) support binding to both AuxRE motifs in the *StPRX28* promoter; the predicted AuxRE-contact surfaces of both ARFs show strongly positive electrostatic potential, mainly due to arginine enrichment (Fig. S12c).

To experimentally confirm the ability of StARF10 and StARF17 to activate the transcription of *StPRX28* gene, we conducted transactivation assays (Lasierra & Prat, 2018) using the ∼1-kb *StPRX28* promoter of NT (containing both predicted ARF-binding motifs) with either StARF10 or StARF17. Both ARFs increased promoter activity: StARF10 by ∼2.2-fold compared to basal promoter activity and StARF17 by ∼1.7-fold (Fig. 4a, Fig. S13, Fig. S14, Table S16, Table S17). These results demonstrate that StARF10 and StARF17 independently activate *StPRX28*, revealing a novel regulatory connection between miR160 and StPRX28.

**Fig. 4.**
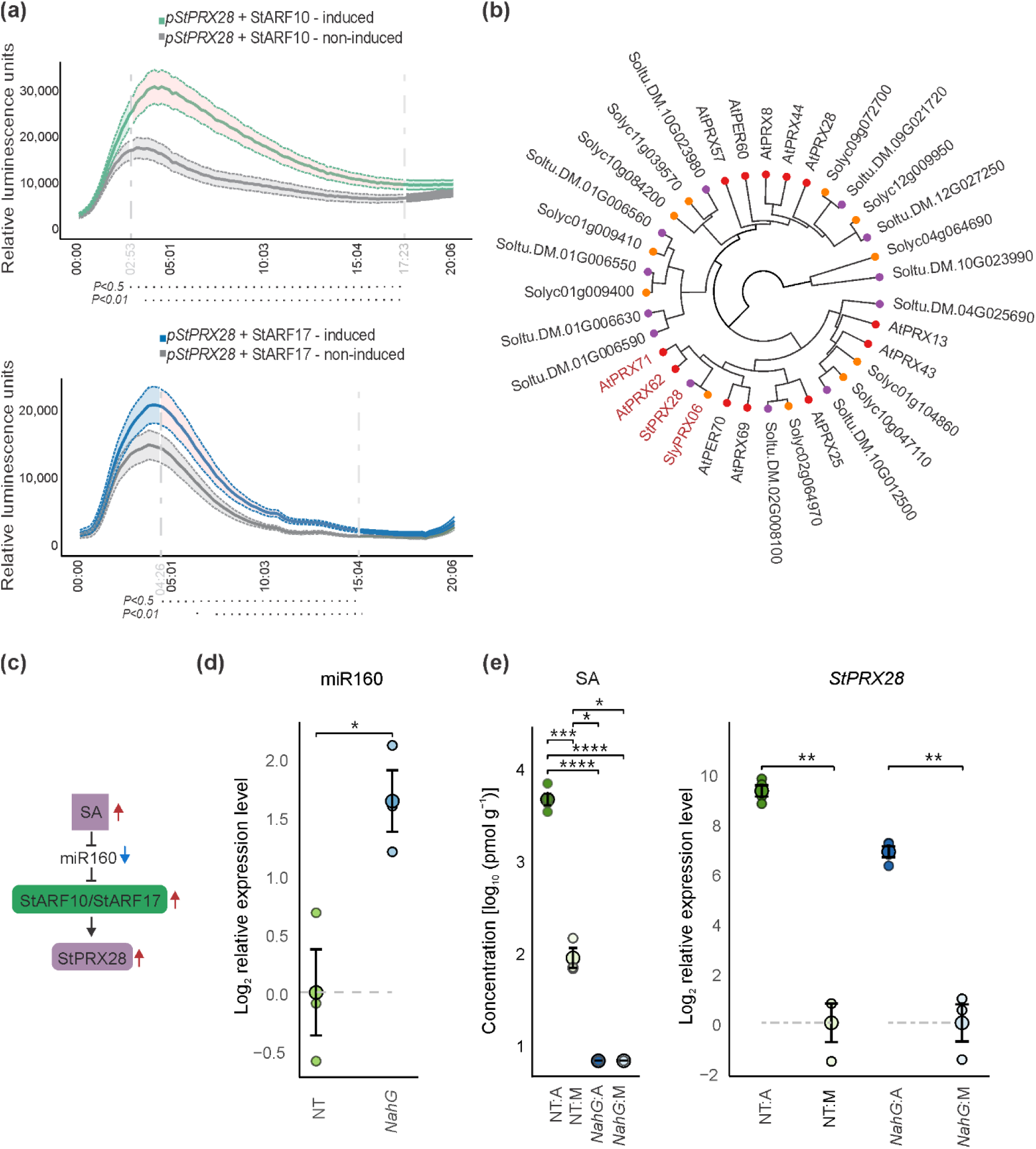
miR160 bridges salicylic acid and ROS signaling. (a) StARF10 and StARF17 activate expression of class III plant peroxidase *StPRX28.* Transactivation assay results showing *in planta StPRX28* promoter (*pStPRX28*) activation by StARF10 and StARF17. The β-estradiol inducible StARF10 (green) and StARF17 (blue) effector vectors and the *StPRX28* promoter (*pStPRX28*) fused to the firefly luciferase reporter were co-infiltrated in *N. benthamiana* leaves. Relative luminescence units (RLU) were measured over time in β-estradiol treated (StARF10 - blue, StARF17 - green) and untreated control (gray) samples. Statistically significant differences between β-estradiol treated and untreated samples (pointwise group comparisons using pairwise Wilcoxon tests with Holm correction) are highlighted with change in ribbon color (*P* < 0.05) and by dot lines below the graphs. Solid lines within the graphs connect mean values of the treated (StARF10 - blue, StARF17 - green) and untreated samples (grey). Dotted lines indicate the ± standard error of the mean for the treated (StARF10 - blue, StARF17 - green) and untreated (grey) samples. Experiments were repeated three times for each ARF, with similar results (Fig. S13, Table S16, Table S17). We also confirmed the expression of GFP-tagged StARF10 and StARF17 only in β-estradiol treated samples (Fig. S14). Confocal microscopy of β-estradiol treated samples showed strong signal of GFP-tagged StARF10 in the nuclei, whereas GFP-tagged StARF17 was detected in the nuclei and cytoplasm (Fig. S14). (b) Phylogenetic tree of class-III peroxidases from *A. thaliana* (At), potato (Soltu; St) and tomato (Solyc; Sly). Circular dendrogram - Affinity Propagation clustering on Levenshtein distance; point colors depict species (red - Arabidopsis, purple - potato, orange - tomato). StPRX28 and its orthologs from Arabidopsis and tomato are marked with red font. (c) Scheme showing the link between SA and StPRX28 regulation via miR160. Colored arrows indicate changes in component levels during the HR to PVY (red, increased; blue, decreased). (d) Level of miR160 in SA-deficient *NahG* line compared to NT (non-transgenic). Group differences were assessed using Welch’s t-test on log₂-transformed data. For visualization purposes values were scaled to the arithmetic mean of the corresponding control. Biological replicates (small circles) and arithmetic mean (large, dark filled circles) ± standard error of the mean are shown. (e) SA is increased in cells adjacent to cell death zones (A section) compared to mock (M)-inoculated tissue sections of NT plants at 5 days post infection (dpi). *NahG* levels were at the limit of detection. Arithmetic mean ± standard error of the mean is shown. Asterisks denote statistically significant differences based on Games-Howell post-hoc test applied to log₁₀-transformed values. *StPRX28* induction level is attenuated in *NahG* relative to NT in A section. Group differences were assessed using Welch’s t-test on log₂-transformed data. For visualization purposes values were scaled to the arithmetic mean of the corresponding control. Biological replicates (small circles) and arithmetic mean (large, dark filled circles) ± standard error of the mean are shown. Asterisks for miR160*, StPRX28* and SA plots denote statistically significant differences (\**P*<0.05, \*\**P*<0.01, *** *P*< 0.001).

As *StPRX28* is a miR160 target strongly induced in our HR system (Fig. 3c), we investigated its functional context through identified orthologs in *Solanum lycopersicu*m and *Arabidopsis thaliana*: SlyPRX06 (RedoxiBase ID: 58), and AthPRX62 and AthPRX71, belonging to the class III peroxidase family (Fig. 4b, Fig S15a). StPRX28 contains heme- and calcium-binding sites, and a signal peptide (Sec/SPI), and it is predicted to localize to the extracellular space (Fig. S15b,c,d), as is typical for Class III peroxidases (Almagro *et al*., 2009; Kidwai *et al*., 2020; Jeong *et al*., 2022). Class III peroxidases regulate apoplastic ROS dynamics, promoting cell-wall cross-linking and stiffening, which limits cell expansion and growth (Almagro *et al*., 2009; Cosio & Dunand, 2009). Both AtPRX71 and AtPRX62 respond to a range of biotic and abiotic stresses—including pathogen infection, wounding, and heavy-metal exposure—and contribute to enhanced resistance against fungal pathogens (Cosio & Dunand, 2009). Presence of multiple *cis*-elements for AP2/ERF, WRKY and TGA/bZIP transcription factors in the *StPRX28* promoter suggests association with ethylene and SA signaling pathways. Given the numerous predicted binding sites for development-associated transcription factors as well (Fig. S12a, Table S12b), *StPRX28* thus likely participates in the coordinated regulation of stress and developmental processes.

### miR160 links SA, auxin and ROS signaling

Given that miR160 promotes auxin-related responses, we asked whether miR160 deregulation perturbs auxin and stress-related hormone balance. We therefore measured auxin/indole-3-acetic acid (IAA), SA, jasmonic acid (JA), and abscisic acid (ABA) levels in fully developed lesions and their surroundings at 5 dpi. Overexpression of miR160 had no detectable effect relative to NT, whereas miR160 depletion resulted in reduced IAA, SA, JA and ABA levels, indicating disrupted hormonal homeostasis and signaling under strong miR160 deficiency (Fig. S16, Table S18). This pattern aligns with extensive phenotypic changes in growth and development of the *mir160a* mutants (Fig. 2a, Fig. S8), which arise primarily from auxin depletion. Elevated miR160 levels in response to PVY in SA-depleted plants (Fig. 1e), and on the other hand strong SA accumulation and concomitantly reduced miR160 levels in NT plants (Fig. 1e, Fig. 4e, Table S18), suggests that SA may act upstream of miR160. To test this, we measured miR160 levels in *NahG* and NT plants under normal growth conditions, to avoid confounding effect of PVY on miR160 expression. Indeed, miR160 levels were increased ∼ 3-fold in *NahG* leaves compared to NT (Fig. 4d, Table S19), supporting SA’s negative regulatory role on miR160. In *NahG*, *StPRX28* was also induced in response to PVY; however, the induction was attenuated relative to NT, indicating fine-tuned miR160-mediated regulation of *StPRX28* (Fig. 4e, Table S20).

Next, we analyzed the promoter regions of *MIR160a* and *MIR160b* genes (∼ 2 kbp) for SA-responsive transcription factor binding sites. We detected multiple binding motifs for bZIP/TGA and WRKY transcription factors in both *MIR160a* and *MIR160b* promoters, and both were also enriched for AP2/ERF binding motifs (Fig. S17, Table S12b). Notably, we also detected ARF-binding motifs in both *MIR160a* and *MIR160b* promoters suggesting a feedback between the miR160 expression and ARF-mediated signaling (Table S12b). Together with previous evidence that miR160 modulates SA accumulation and signaling (Natarajan *et al*., 2018) our results point to a bidirectional SA-miR160 circuit and place SA upstream of StARF10 and StARF17 within SA–auxin crosstalk.

### miR160 increases sensitivity to heat in potato

Given that miR160 OE plants exhibited leaf epinasty, whereas *miR160b*-deficient mutants showed hyponastic movement of leaves (Fig. 2a)—a phenotype reminiscent of heat stress symptoms in potato (Zagorščak *et al*., 2025)—we investigated whether these miR160-mediated morphological changes would confer distinct adaptive outcomes at elevated temperature. Under heat stress (30 °C day/28 °C night), NT and miR160 OE lines developed necrosis and leaf yellowing, with lower leaves being lost earlier in miR160 OE lines than in NT. By contrast, *mir160a* and *mir160b* mutants remained phenotypically unaffected under heat stress (Fig. 5a, Fig. S18, Fig. S19). Unlike NT and OE lines, both miR160-depleted lines maintained stable photosynthetic performance and leaf water status (as indicated by a lower water index) (Fig. 5b, Table S21). A functional link between the miR160–ARF10 module and stomatal-mediated water regulation has been suggested, as ARF10 was reported to promote development of larger stomata and to increase transpirational water loss in tomato (Liu *et al*., 2016). Enlarged stomata under heat stress, however, may be favorable for increasing transpirational water loss and enhancing evaporative cooling, thereby protecting leaves from heat-induced damage. *mir160a* mutants develop larger guard cells and stomata than NT plants (Fig. 2c, Fig. S7). However, despite this increase in size, leaf perspiration remained similar under heat stress for all analyzed genotypes (Table S21). Consistently, ΔT measurements, functional proxy for stomatal conductance and evaporative cooling, also did not differ among genotypes (Table S21). Together, these results indicate that the heat-tolerance phenotype cannot be explained solely by stomatal enlargement and likely involves additional mechanisms.

**Fig. 5:**
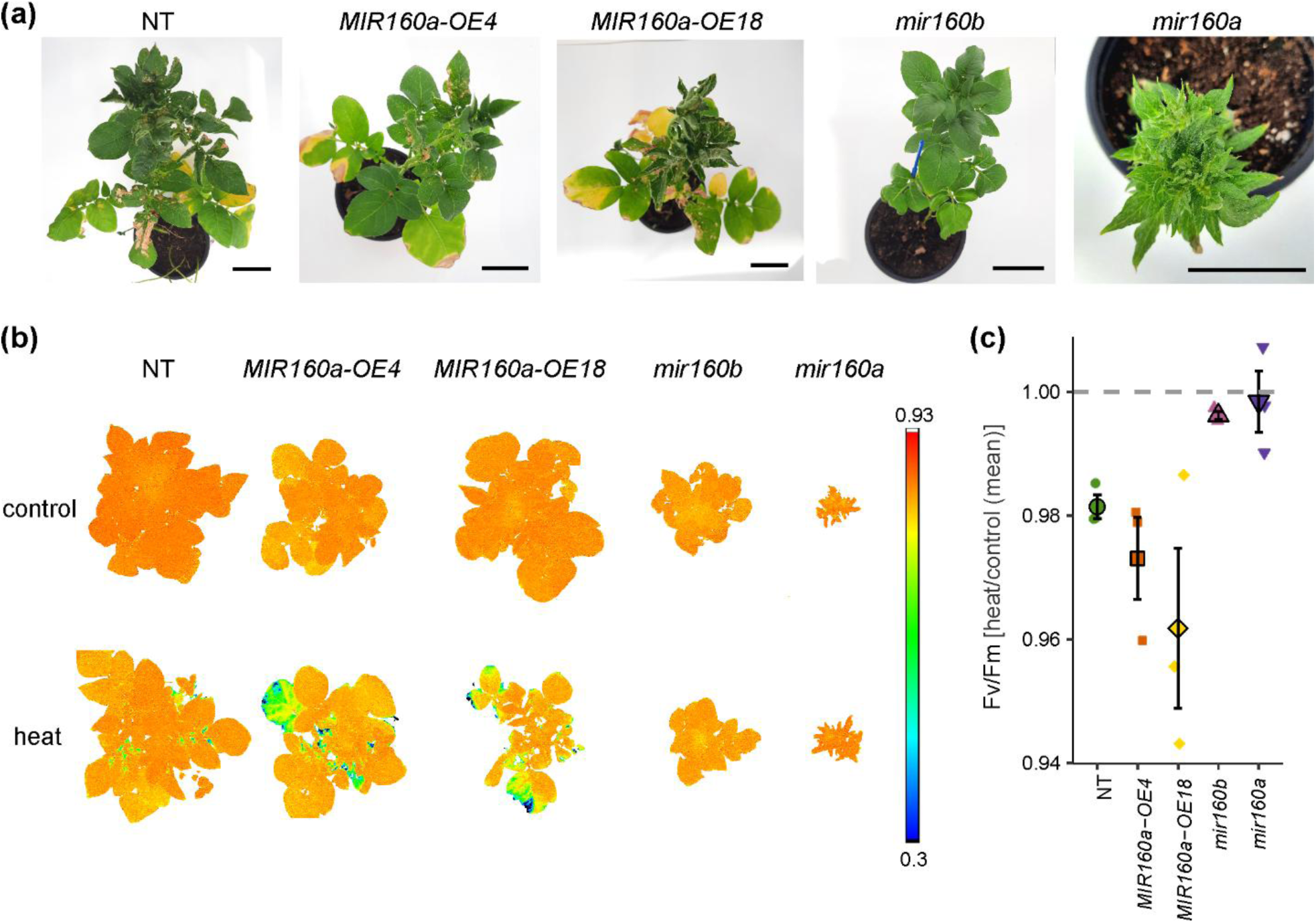
miR160 increases potato susceptibility to heat stress. (a) Heat-stress phenotypes of miR160 overexpression lines (*MIR160a-OE4*, *MIR160a-OE18*), miR160-depleted lines (*mir160b*, *mir160a*), and NT (non-transgenic) controls after 15 days of heat stress at 30/28 °C (day/night). NT and miR160 OE plants show heat-induced necrosis and leaf yellowing/senescence, whereas miR160-depleted plants remain symptomless. The top view of the plants is shown, for the side view see Fig. S18 and Fig. S19. Scale bar: 1cm. (b) False-color chlorophyll-fluorescence images of the maximal quantum efficiency of photosystem II (Fv/Fm) acquired with high-throughput phenotyping PlantScreen™ System after dark adaptation after 10 days of heat stress. Color scale bar represents the range of fluorescence values. (c) Maximum quantum efficiency of photosystem II measures in potato plants after 10 days of heat stress, normalized to non-heat-treated controls. Maximum quantum efficiency of photosystem II reflects the photosynthetic efficiency of the plants. For visualization purposes, heat measurements were scaled to the arithmetic mean of the corresponding controls.

## Discussion

The hypersensitive response (HR) has been described over a century ago, yet mechanistic basis of viral arrest is still not fully understood. While transcriptional regulation of HR has been studied to some degree, the contribution of post-transcriptional, sRNA–mediated processes has remained unexplored.

Here, using spatially resolved sRNAomics approach, we found that miR160 is specifically downregulated in cells adjacent to cell death zones of Ny-1 dependent HR, but not in sensitive SA-depleted genotype, (Fig. 1e, Table S6). Notably, previous studies mostly reported miR160 induction across diverse plant-pathogen interactions—often in PTI or compatible infection contexts (Li *et al*., 2010; Du *et al*., 2011; Yin *et al*., 2013). However, broader functional role of miR160 in stress and development has not yet been investigated.

Our functional characterization confirmed that in the absence of stress, miR160 is indispensable for coordinated growth and development in potato. Reducing miR160 produced graded outcomes: partial depletion caused leaf hyponasty, whereas stronger depletion resulted in severe growth defects, revealing a dosage-sensitive response (Fig. 2a, Fig. S8). Post-transcriptional regulation of Class C ARFs, *StARF10* and *StARF17* by miR160 is well-established. Here we uncovered a broader miR160 impact, spanning auxin signaling, cell-wall dynamics, ROS and heat shock stress pathways (Fig. 2d,e, Fig. 3b,c, Table S10, Table S11). Importantly, we also link miR160 to tuberization: miR160 negatively regulates three well-known repressors of tuberization *StFKF1*, *StGI* (GIGANTEA), and *StSPG5* (Fig. 2e, Fig. 3b, Table S10, Table S14) (Salaria *et al*., 2020; Bao *et al*., 2025). Such repression may facilitate tuberization under normal growth conditions. In line with this, miR160 depletion is associated with diminished tuberization (Fig. 2a, Fig. S8, Table S9).

Overaccumulation of miR160, however, promoted susceptibility-like traits—leaf epinasty and senescence (Fig. 2a, Fig. 5a, Fig. S9, Fig. S18, Fig. S19, Fig. S20). Leaf epinasty is a common disease symptom across several host-virus interactions (e.g. begomoviruses, CMV, RSMV, TMV) (Gonsalves *et al*., 1994; Song *et al*., 2009; Li *et al*., 2020a; Dai *et al*., 2022; Ma *et al*., 2023; Jiang & Zhou, 2023) and is also observed in the PVY-*N.benthamiana* pathosystem (Fig. S21). This aligns with evidence that plant viruses frequently manipulate auxin signaling pathway to facilitate infection (Zhang *et al*., 2020). Supporting this, ARF17, a common target of different viral proteins, acts as a positive regulator of antiviral resistance, and its depletion correlates with more severe disease symptoms (Zhang *et al*., 2020; Ma *et al*., 2023). We here found that in sensitive SA-depleted *NahG* plants, PVY infection increased miR160 accumulation (Fig. 1e), pointing to PVY interference with miR160 pathway upstream of ARFs. Indeed, it is known that viral-encoded silencing suppressors, including potyviral HC-Pro, can interfere with miRNA-guided regulation and induce developmental abnormalities (Kasschau *et al*., 2003; Jin *et al*., 2025), suggesting that perturbed miR160-directed ARF regulation contributes to symptom development. Despite PVY-driven miR160 induction in *NahG*, leaf epinasty did not develop, likely because the magnitude of induction (∼3 fold) was far lower than that of miR160 overexpression plants (∼30 fold). However, PVY-infected *NahG* plants displayed enhanced senescence relative to NT at later time points after infection (Fig. S2).

Crosstalk between SA and auxin has been proposed downstream of ARFs at the level of GRETCHEN HAGEN 3.6, an auxin/SA adenylating enzyme (Natarajan *et al*., 2018). Our data position miR160 at the SA-auxin interface, acting as a hub that coordinates defense–growth outputs. SA is central to immunity against biotrophic pathogens, including viruses, and SA accumulation at infection sites is a hallmark of ETI (Betsuyaku *et al*., 2018; Radojičić *et al*., 2018; Zhang & Li, 2019). Accordingly, in our viral HR system SA was elevated in cells adjacent to the cell-death zone (Fig. 4e, Table S18). Higher SA coincided with reduced miR160 levels (Fig. 1e), indicating a rebalancing of the network from growth/development regulation toward defense activation. Consistent with this, multiple miR160-repressed target genes, were strongly induced during PVY-triggered HR—particularly in the immediate surroundings of the cell death zone—including genes involved in cellular energy metabolism, repressors of tuberization, and regulators of ROS homeostasis (Fig. 3b,c, Table S14). A link between miR160 and ROS accumulation (H₂O₂) has already been reported in fungal pathosystem (Li *et al*., 2014); however, the signaling circuit connecting miR160 and ROS remained so far unresolved. In this study, we identify a miR160–StARF10/StARF17–StPRX28 module as one mechanistic route coupling miR160 and ROS signaling pathway (Fig. 4).

StPRX28 belongs to Class III apoplastic peroxidases which form a complex family of proteins that catalyze the oxidoreduction of various substrates using H_2_O_2_. In particular, pH-dependent peroxidases in the cell wall can also be a source of apoplastic H₂O₂ (Torres *et al*., 2006). Thus, peroxidases have dual roles in both generation and scavenging of ROS. This function enables them to fine-tune redox dynamics to sustain effective defense signaling while preventing oxidative stress-induced cellular damage (Minibayeva *et al*., 2015; Kidwai *et al*., 2020). The expression of peroxidase enzymes has been shown to be induced following recognition of bacterial and fungal pathogens (Chittoor *et al*., 1997; Sasaki *et al*., 2004; Torres *et al*., 2006). Consistently, we found that *StPRX28* is strongly induced in ETI of potato to PVY infection (Fig. 3c, Table S15). On the other hand, evidence from several systems links diminished activity or expression of Class III peroxidase with impaired oxidative burst capacity and increased pathogen susceptibility (Bindschedler *et al*., 2006; O’Brien *et al*., 2012). In *A. thaliana*, cell wall peroxidases such as PRX33 and PRX34 promote apoplastic ROS accumulation and cross-linking of cell wall polysaccharides upon recognition of diverse MAMPs (Bindschedler *et al*., 2006; Daudi *et al*., 2012; O’Brien *et al*., 2012). In pepper, the Class III peroxidase PRX02 was shown to be essential for HR elicited by *Xanthomonas campestris* pv. *vesicatoria* (Choi *et al*., 2007). Similarly, PRX25 from *Citrus sinensis* has been implicated in fine-tuning ROS to promote stronger yet more controlled HR, enhancing disease resistance, without causing excessive oxidative damage (Li *et al*., 2020b). Most interestingly, expression of AtPRX71—ortholog of StPRX28 (Fig. 4b)—restricts growth while promoting ROS accumulation, lignification, and cell-wall strengthening (Raggi *et al*., 2015). This phenotype exemplifies a classic growth–defense trade-off: ROS-driven cross-linking shifts carbon usage toward barrier construction, thereby prioritizing defense over growth.

Previous studies have shown that SA can stimulate ROS production through both cytosolic and apoplastic peroxidases, while ROS in turn enhance SA accumulation, forming a positive feedback loop that amplifies plant defense responses (Herrera-Vásquez *et al*., 2015; Lukan *et al*., 2020). We propose a model (Fig. 6) in which SA represses miR160, thereby alleviating StARF10/StARF17-mediated repression of *StPRX28* in cells adjacent to cell death zones. In parallel, transcriptional regulation by SA-responsive transcription factors (e.g., WRKY/TGA) may also enhance *StPRX28* transcription. Such dual modes of control likely enhance the robustness of the immune response (Tsuda *et al*., 2013). Notably, presence of enriched AP2/ERF motifs in the *StPRX28* promoter suggests modulation by the ethylene branch as well, consistent with reports that many peroxidases respond to SA and ethylene (El-Sayed & Verpoorte, 2004).

**Fig. 6.**
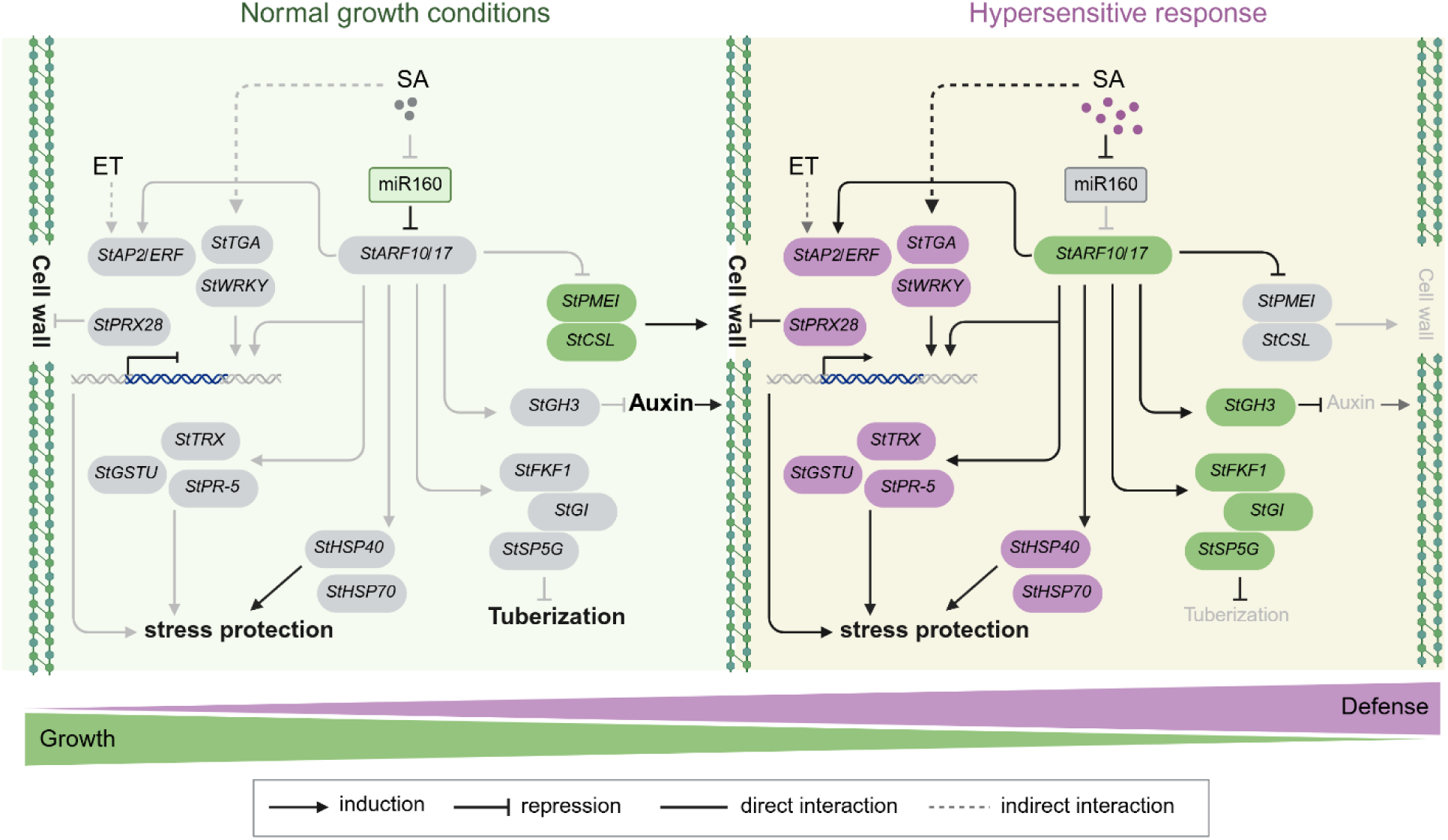
Current knowledge-based model of the miR160-mediated regulatory network. Under normal growth conditions (left), miR160 promotes growth and development. In hypersensitive response to PVY (right), salicylic acid (SA) levels increase and miR160 is repressed. This derepresses *StARF10* and *StARF17* and activates downstream signaling, including genes involved in ROS homeostasis and stress protection, prioritizing defense-related outputs at the cost of growth, indicative of a growth–defense trade-off. In each condition, active modules and interactions (induction, repression) are color-coded as growth (green) or defense (purple), active interactions are shown in black, whereas inactive modules and interactions are shown in grey. Created in BioRender.com.

Furthermore, miR160 represses *StHsp70* and *StHsp40* known to promote thermotolerance (Nover & Scharf, 1997; Kotak *et al*., 2007; Wang *et al*., 2021). We show that miR160 promotes susceptibility to heat stress, as both ∼50% and ∼80% depletion enhance tolerance to heat stress. Importantly, partial (∼50%) depletion provides a particularly advantageous balance, substantially improving stress resilience while exerting minimal effects on potato growth and tuberization (Fig. 2a,c, Fig. 5, Fig. S8, Fig. S9, Fig. S18, Fig. S19, Table S9).

Increasing stress resilience and yield are central breeding goals, yet maximizing growth-related traits can inadvertently lead to the loss of useful genetic traits for defense (He *et al*., 2022). Thus, balancing growth and plant defense is crucial for crop varieties that couple high yield with robust resilience. Our work identifies miR160 as a signaling hub linking SA, auxin, heat shock and ROS pathways and thereby modulating growth and responses to virus infection and heat stress. Its dual role—as a negative regulator of HR-associated genes and a negative regulator of thermotolerance—positions miR160 as a promising target for precision modulation, where partial repression could deliver multi-stress tolerance with minimal yield loss.

## Acknowledgements

The authors thank Tjaša Mahkovec Povalej, Barbara Jaklič, Tina Arnšek, Rebeka Strah, Anja Moškrič for technical support and laboratory assistance. This research was financially supported by the Slovenian Research and Innovation Agency grants P4-0165, P4-0463, J4-2544, and Czech Science Foundation grant 22-17435S. The authors also acknowledge the Slovenian Research Agency for supporting research infrastructure used in this work through grant IO-0004, the Slovenian node of Euro-BioImaging ERIC and the Slovenian node of Instruct ERIC for providing access to imaging technologies.

## Competing interests

None declared.

## Author contributions

MK, TL, AC, KG planned and designed the research. MK, KP, KS, NM, VL, JS, ON, AK performed the experiments. MK, MZ, KPO, KS, VL, JS, ON analyzed the data. All authors contributed to the writing or revision of the article.

## Data availability

The sRNA-Seq and RNA-seq raw Illumina reads have been deposited in European Nucleotide Archive under accession numbers PRJEB108023 and PRJEB108039. All data analysis scripts are publicly available in the GitHub repository at https://github.com/NIB-SI/miR160-HR. Microscopy datasets are available on Zenodo (10.5281/zenodo.18797111).

## Supporting Information (brief legends)

Fig. S1: Diversity of CRISPR-Cas9-induced mutations at the *MIR160b* locus in stably transformed *mir160b* plants in the cv. Rywal genetic background.

Fig. S2: SA-depleted *NahG* plants show impaired PVY resistance with systemic lesion development.

Fig. S3: sRNA response is diminished beyond the lesion margins.

Fig. S4: PVY spread around the cell death zone in *NahG*.

Fig. S5: Comparison of pre-miR160a and pre-miR160b precursor sequences and their predicted secondary structures.

Fig. S6: Reduced miR160 levels affects mesophyll organization.

Fig. S7: Strongly reduced miR160 levels results in development of larger guard cells.

Fig. S8: Strong miR160 depletion diminishes root growth and tuber formation.

Fig. S9: Reduced miR160 levels are associated with delayed senescence.

Fig. S10: PVY remains confined to the cell death zone in miR160 overexpression plants.

Fig. S11: miR160 represses *StHSP70* and *StPRX28*.

Fig. S12: StARF10 and StARF17 recognize AuxRE in *StPRX28* promoter.

Fig. S13: StARF10 and StARF17 activate expression of class III plant peroxidase *StPRX28*.

Fig. S14: Confirmation of expression of GFP-tagged StARF10 and StARF17 under β-estradiol inducible promoter in *N. benthamiana* leaves.

Fig. S15: StPRX28 is Class III apoplastic peroxidase.

Fig. S16: Hormone levels in fully developed lesions and surrounding tissue at 5 days post infection (dpi) in miR160-overexpression (*MIR160-OE4 and MIR160-OE18*) and miR160 knockdown (*mir160a*) lines, relative to non-transgenic (NT) plants.

Fig. S17: Enriched transcription-factor binding motifs in the *MIR160a* and *MIR160b* promoters.

Fig. S18: miR160 increases sensitivity to heat in potato (Experiment No. 1).

Fig. S19: miR160 increases sensitivity to heat in potato (Experiment No. 2).

Fig. S20: Increased miR160 levels are associated with accelerated senescence at later time points.

Fig. S21: PVY induces leaf curling in *Nicotiana benthamiana*.

Table S1: Plasmid sequences from positive colonies of overexpression constructs and diversity of CRISPR-Cas9-induced mutations at the *MIR160b* locus in stably transformed *mir160b* plants in the cv. Rywal genetic background.

Table S2: Selected miRNAs and genes with corresponding primers and probes sequences for expression analyses with quantitative PCR.

Table S3: Oligonucleotides used for cloning for transactivation assay.

Table S4: Spatial regulation of sRNAs in lesion and immediately adjacent cells, and the surrounding regions of resistant Rywal (NT) and susceptible *NahG-Rywal* (NahG) plants at 3 days post infection (dpi).

Table S5: Differentially expressed sRNAs and their target levels from sRNA-seq and RNA-seq at 3 days post inoculation (dpi).

Table S6: miR160 shows differential spatial regulation in cells adjacent to the cell death zone between NT and *NahG* plants.

Table S7: Levels of miR160, *StARF10*, and *StARF17* in miR160 overexpression and knockdown plants relative to cv. Rywal (NT).

Table S8: Guard cell size of miR160 overexpression and knockdown plants relative to cv. Rywal (NT).

Table S9: Strong miR160 depletion impairs tuberization.

Table S10: Effect of miR160 deregulation on gene expression under normal growth conditions.

Table S11: Gene set enrichment analysis (GSEA) results.

Table S12: Transcription factor binding motifs in the promoter region of selected genes and miRNAs.

Table S13: Relative RNA abundance in systemic leaves 18 days post infection (dpi).

Table S14: Gene expression changes in lesion and immediately adjacent cells, and the surrounding regions at 5 days post infection (dpi) following PVY infection.

Table S15: miR160 negatively regulates expression of *StHSP70* and *StPRX28*.

Table S16: Relative luminescence (LUM) measurements showing StARF10-mediated activation of the class III plant peroxidase *StPRX28*.

Table S17: Relative luminescence (LUM) measurements showing StARF17-mediated activation of the class III plant peroxidase *StPRX28*.

Table S18: Hormone levels in overexpression (*MIR160a-OE4, MIR160a-OE18*), knockdown (*mir160a*) and *NahG-Rywal* (NahG) plants relative to cv. Rywal (NT) at 5 days post infection (dpi).

Table S19: miR160 levels are increased in *NahG-Rywal (NahG)* plants compared with cv. Rywal (NT) plants.

Table S20: *StPRX28* induction level is attenuated in *NahG-Rywal (NahG)* relative to cv. Rywal (NT) in the cells adjacent to cell death zones.

Table S21: miR160-depleted lines maintained stable photosynthetic performance and leaf water status.

Dataset S1: Dataset S1. sRNA–target regulatory network connecting differentially expressed sRNAs and their predicted target transcripts at 3 days post inoculation (3 dpi). For each predicted interaction, the inferred mode of regulation (cleavage or translational repression) is provided based on *in silico* prediction using psRNATarget_v2 (Cleavage_in_silico; Translation_repression_in_silico) and degradome evidence (Cleavage_PARE). Node colors indicate expression changes: red, upregulated; blue, downregulated; grey, not significant (FDR-adjusted *P*-value < 0.05).

Dataset S2. miR160-target regulatory network. Network was constructed from genes showing significant differential expression in miR160 overexpression and knockdown plants compared with non-transgenic (NT) plants under normal and PVY-infected conditions. Node colors indicate expression changes: red-upregulated, blue-downregulated. miR160 direct targets are labeled plotRank1; miR160-regulated genes with ARF-binding sites identified in their promoter regions are labeled plotRank2; miR160-regulated genes without detectable ARF-binding sites are labeled plotRank3.

## Notes

### Competing Interest Statement

The authors have declared no competing interest.

