## Supplementary for "miR160 controls class III peroxidase via *StARF10/17* and couples salicylic acid and ROS signaling networks in potato hypersensitive response to potato virus Y"

**(a)**

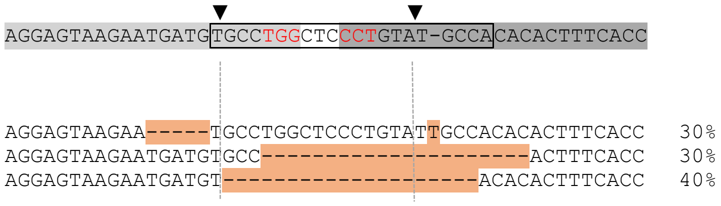


**(b)**

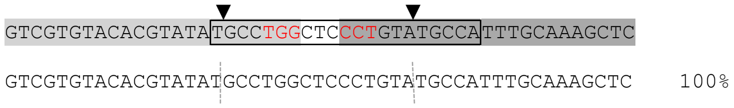


**Fig. S1: Diversity of CRISPR-Cas9-induced mutations at the *MIR160b* locus in stably transformed *mir160b* plants in the cv. Rywal genetic background**. Results of sequencing for individual PCR products within (a) *MIR160b* locus where we expected mutations and as a control within (b) *MIR160a* locus to exclude the possibility of off-target effects in the homologous region. Next to sequences, the percentage of PCR products with the particular type of mutations is shown. Ten PCR products were sequenced per locus. Mature miR160 (boxed), sgRNA1 (light grey), sgRNA2 (dark grey), PAM motif (red), theoretical cutting site (arrowhead) as located in the precursor are shown above the alignment. Mutations are shown in orange. Primers used for sequencing are specified in Lukan *et al.* (2022). Sequencing data of all colonies for both *MIR160a* and *MIR160b* loci are given in Table S1.

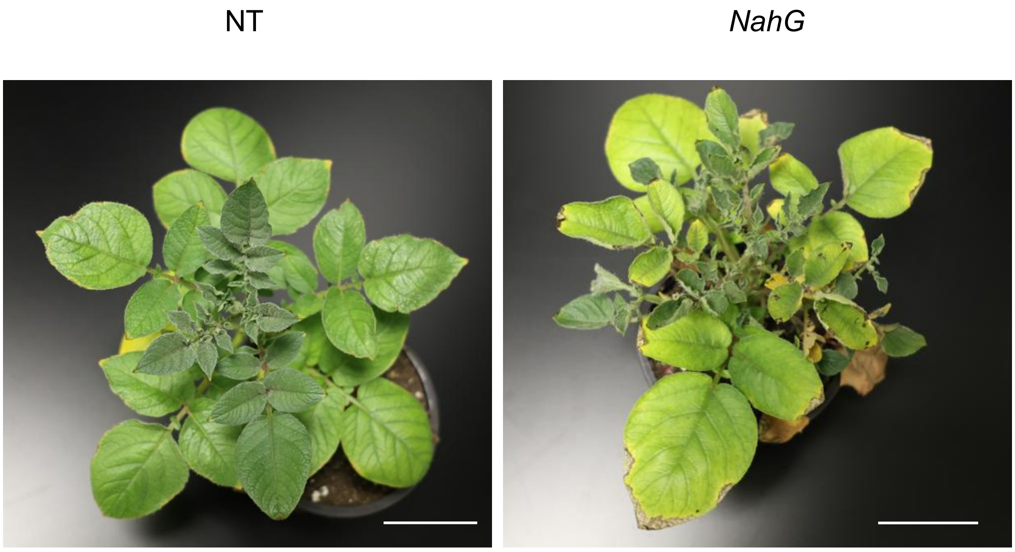


**Fig. S2: SA-depleted *NahG* plants show impaired PVY resistance with systemic lesion development.** Symptom development on upper noninoculated leaves in *NahG* plants at 21 days post inoculation (dpi). Non-transgenic PVY-resistant cv. Rywal (NT) plants at 21 dpi show no symptoms. Scale bar: 5cm.

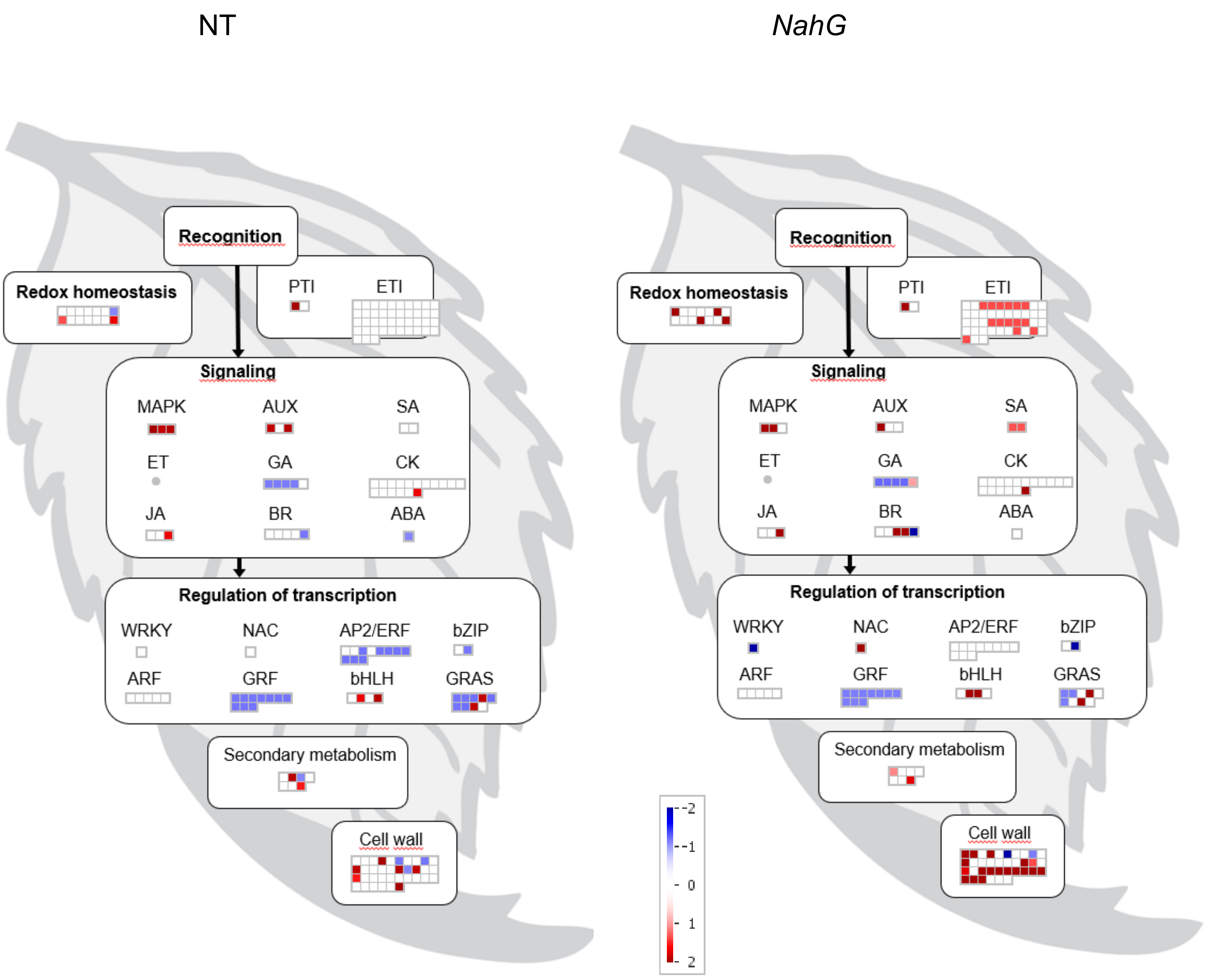


**Fig. S3: sRNA response is diminished beyond the lesion margins.** Visualization of differentially expressed sRNAs identified in the surrounding tissue (section B) of early developed lesions at 3 days post inoculation (dpi) in cv. Rywal (NT) and *NahG*, according to the function of their predicted targets. Each square represents one target gene and colors represent $log_2$ ratios of sRNA expression between PVY- and mock-inoculated plants (red - upregulated; blue - downregulated; FDR-adjusted $p < 0.05$). MapMan4 ontology Bins: Redox homeostasis (10), Pathogen.pattern-triggered immunity (PTI) machinery (26.10.1), Pathogen.effector-triggered immunity (ETI) machinery (26.10.2), MAPK (27.14), AUX (11.2), SA (11.8), ET (11.5), GA (11.6), CK (11.4), JA (11.7), BR (11.3), ABA (11.1), WRKY (15.5.7.5), NAC (15.5.7.1), AP/ERF (15.5.7.6), bZIP (15.5.1.1), ARF (15.5.9.1), GRF (15.5.2.6), bHLH (15.5.1.2), GRAS (15.5.10.2), secondary metabolism (9), cell wall (21).

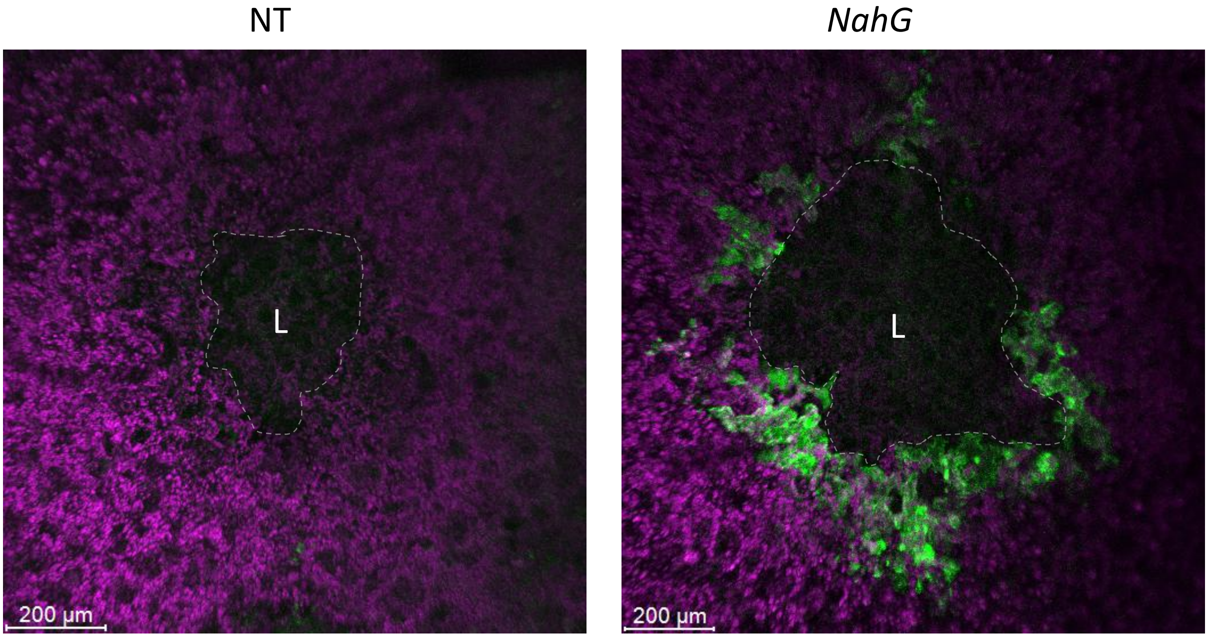


**Fig. S4: PVY spread around the cell death zone in *NahG*.** PVY-N605(123)-GFP accumulation (green) was analyzed at fully developed lesion stage (at 6 days post inoculation) by confocal microscopy. Figures represent the overlay of PVY-N605(123)-GFP accumulation (green) and chlorophyll fluorescence (purple). The lesion is marked by the white "L", and the edge of the cell death zone is marked by the light dotted line. Scale bar: 200 µm. NT: non-transgenic cv. Rywal.

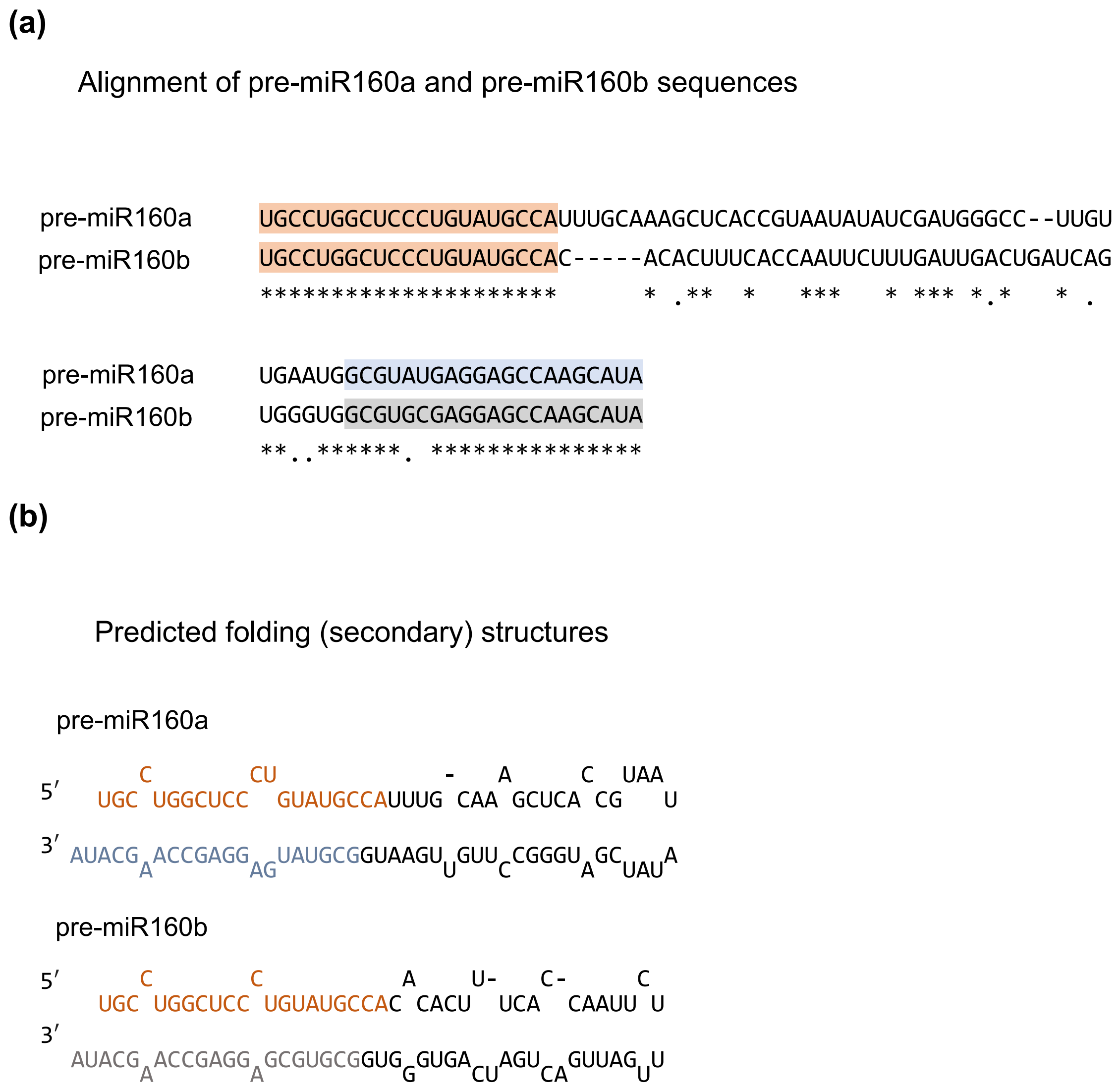


**Fig. S5: Comparison of pre-miR160a and pre-miR160b precursor sequences and their predicted secondary structures.** (a) Alignment of pre-miR160a and pre-miR160b sequences (source: https://www.mirbase.org/). (b) Predicted secondary structure of pre-miR160a and pre-miR160b. Both *MIR160* loci produce identical miR160 from 5´arms of precursors transcripts (miR60a,b-5p; orange) whereas miR160 from 3´ends differ (pre-miR160a precursor miR160a-3p; blue, pre-miR160b precursor produce miR160b-3p; gray).

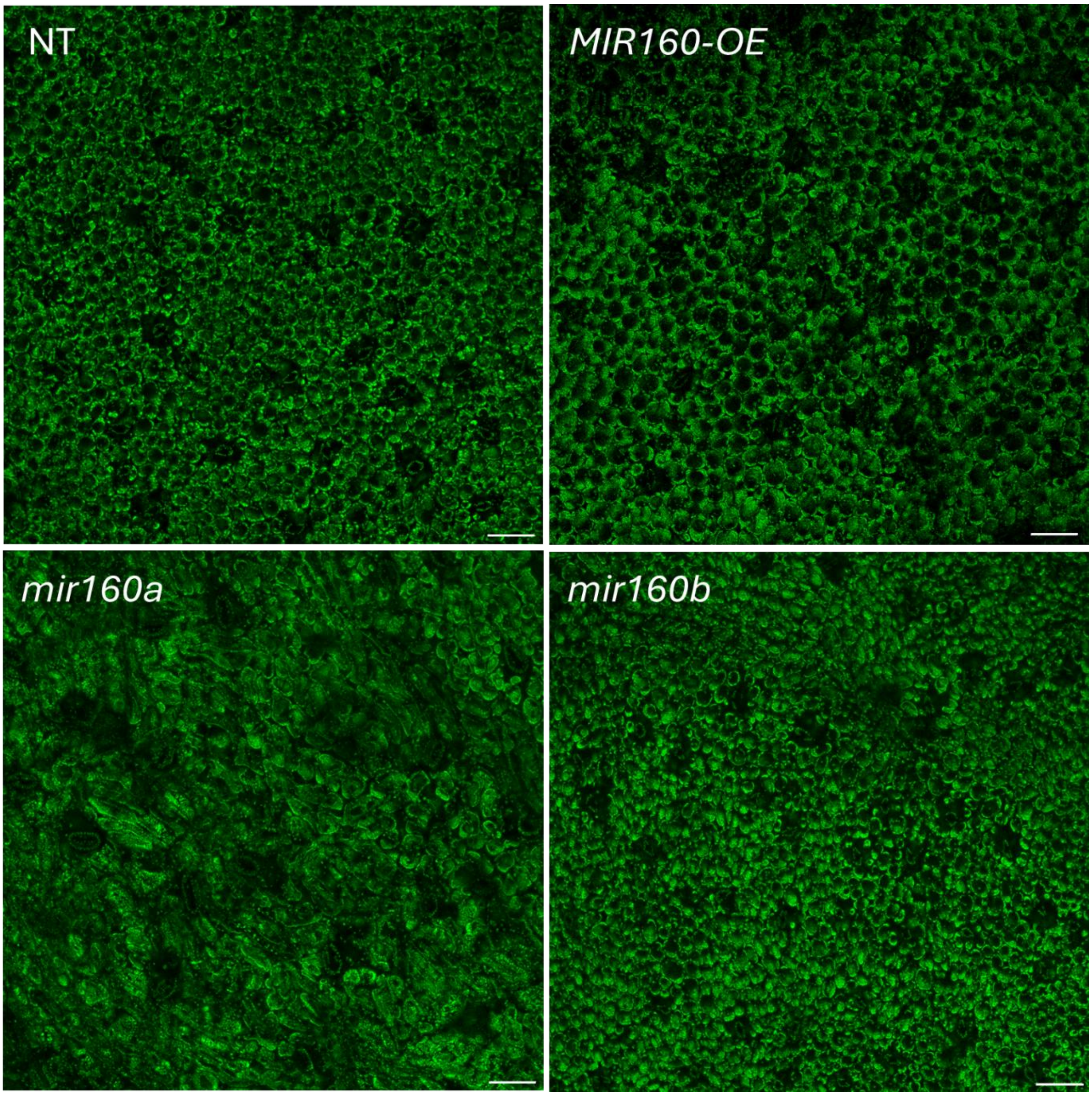


**Fig. S6: Strongly reduced miR160 levels affects mesophyll organization**. Leaf tissue organization of miR160 overexpression (*MIR160-OE4*) and miR160 knockdown lines (*mir160a*, *mir160b*) visualized through chlorophyll autofluorescence in green by confocal microscopy. Only *mir16*0a knockdown lines showed variably shaped cells of palisade tissue. Scale bar: 100 µm. NT: non-transgenic cv. Rywal.

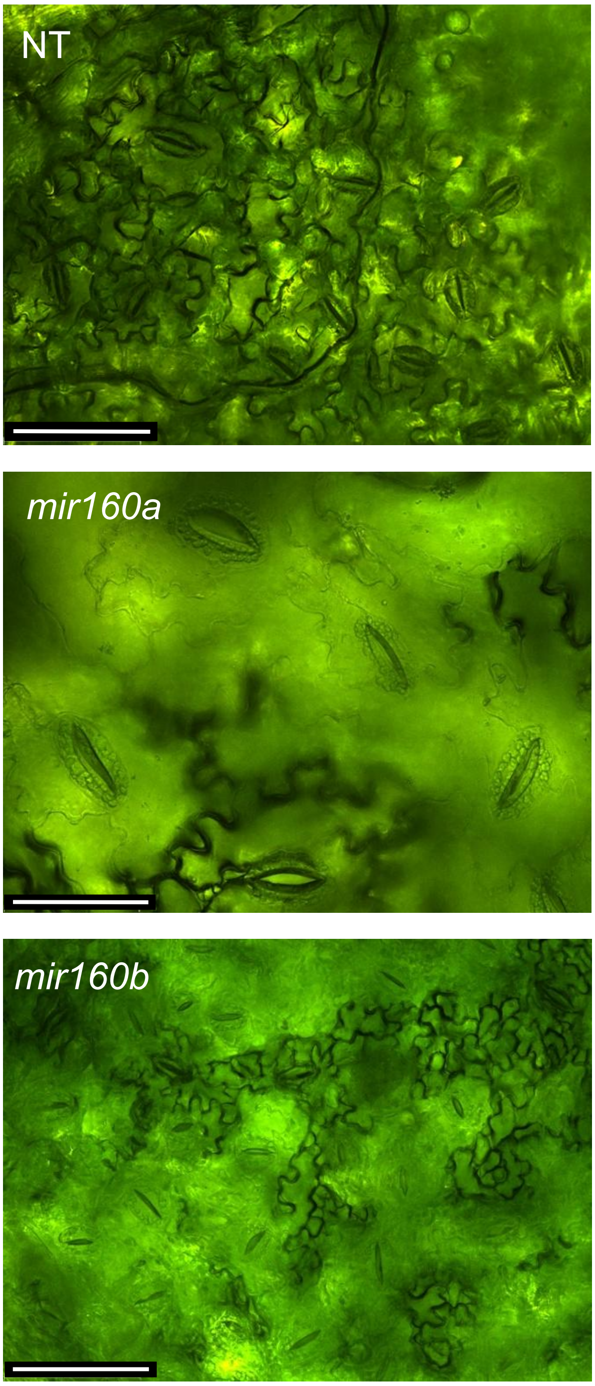


**Fig. S7: Strongly reduced miR160 levels results in development of larger guard cells.** The abaxial leaf surface of cv. Rywal (NT) and miR160 knockdown lines (*mir160a*, *mir160b*) was imaged by brightfield microscopy using a digital camera. Scale bar: 100 µm.

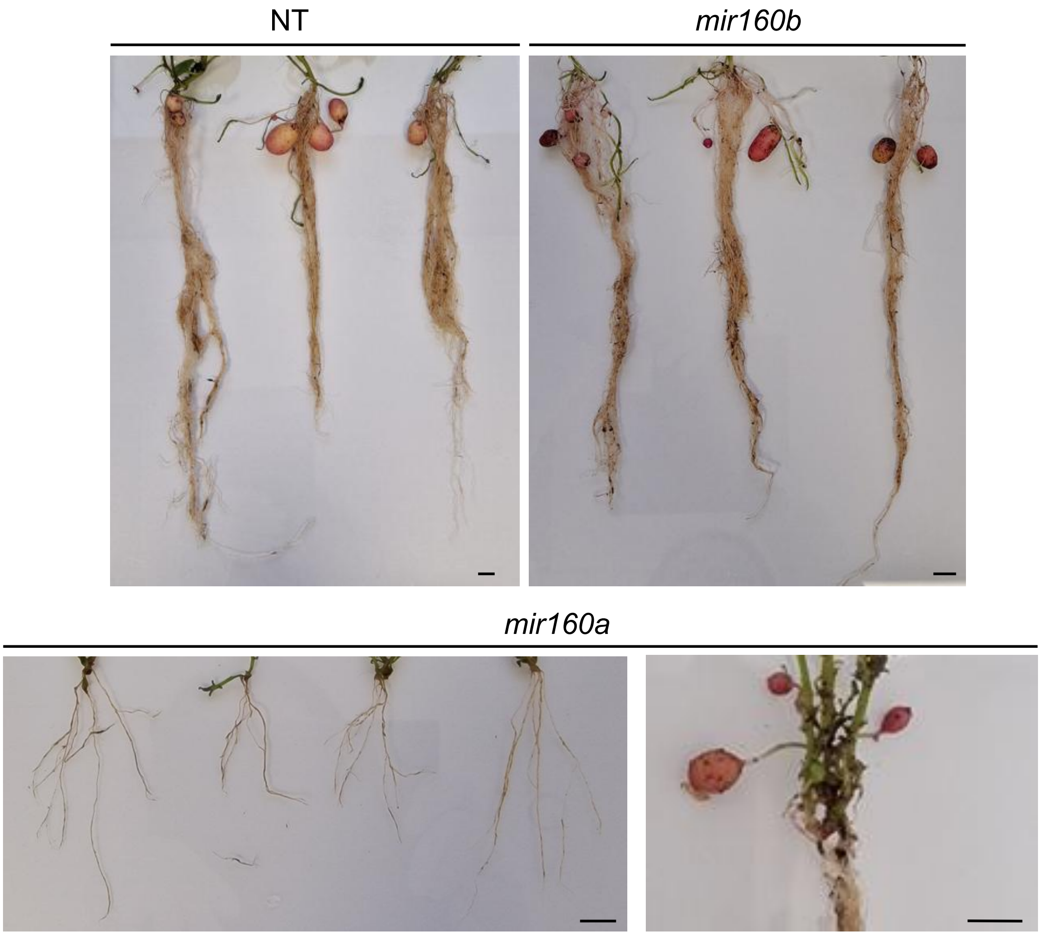


**Fig. S8: Strong miR160 depletion diminishes root growth and tuber formation**. Photographs show roots of 5-week-old NT (non-transgenic) plants compared with *mir160b* and *mir160a* lines. In *mir160a* plants, tubers were not observed at 5 weeks (lower left panel) but developed with a delay and were detected at 12 weeks (lower right panel). Scale bar: 1 cm.

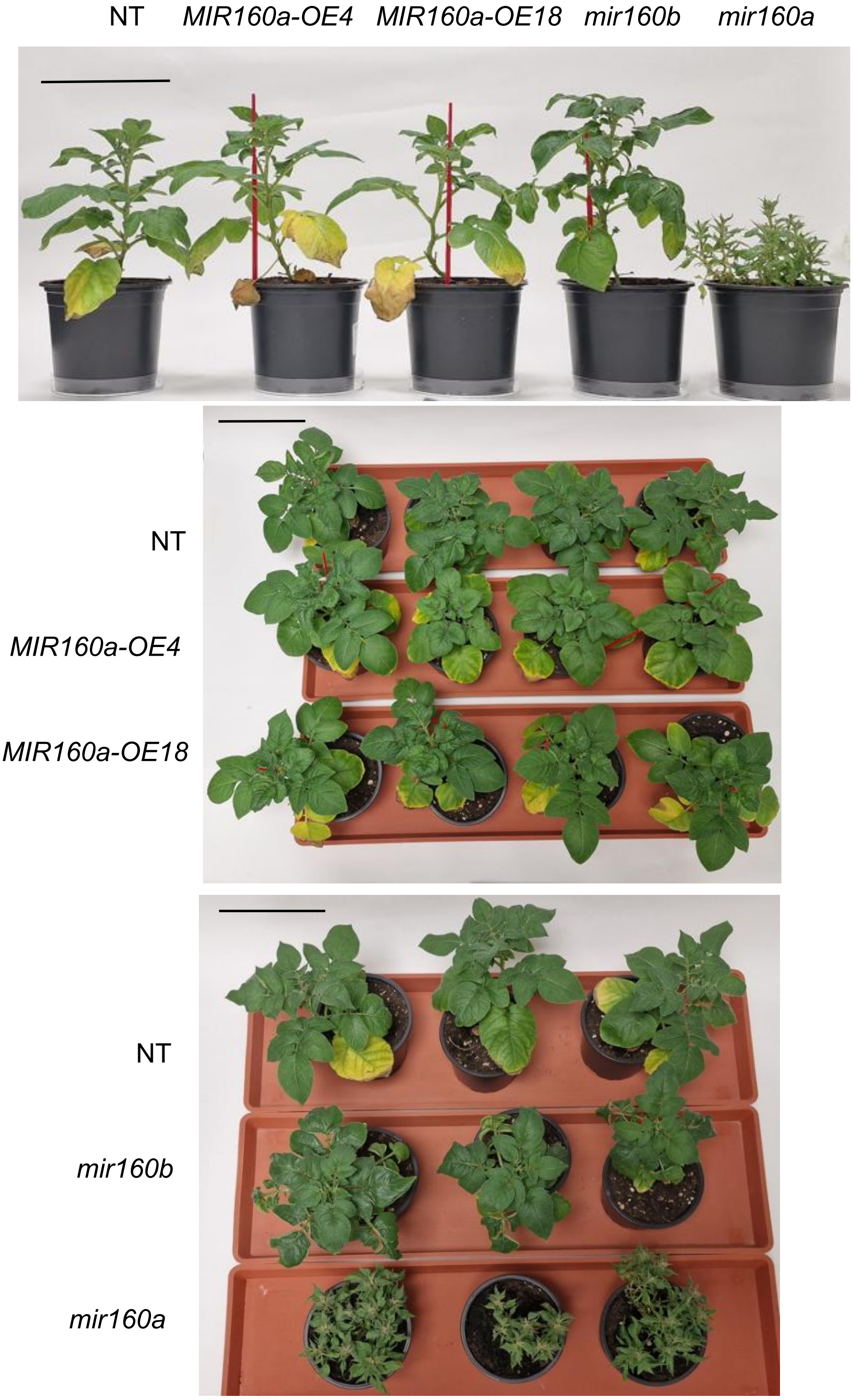


**Fig. S9: Reduced miR160 levels are associated with delayed senescence.** Phenotypes of 5-week-old miR160 overexpression lines (*MIR160a-OE4*, *MIR160a-OE18*) and knockdown lines (*mir160a*, *mir160b*) grown under normal conditions, compared with NT (non-transgenic) potato. The top panels show side views of the plants, and the bottom panels show top views. Scale bar: 10 cm.

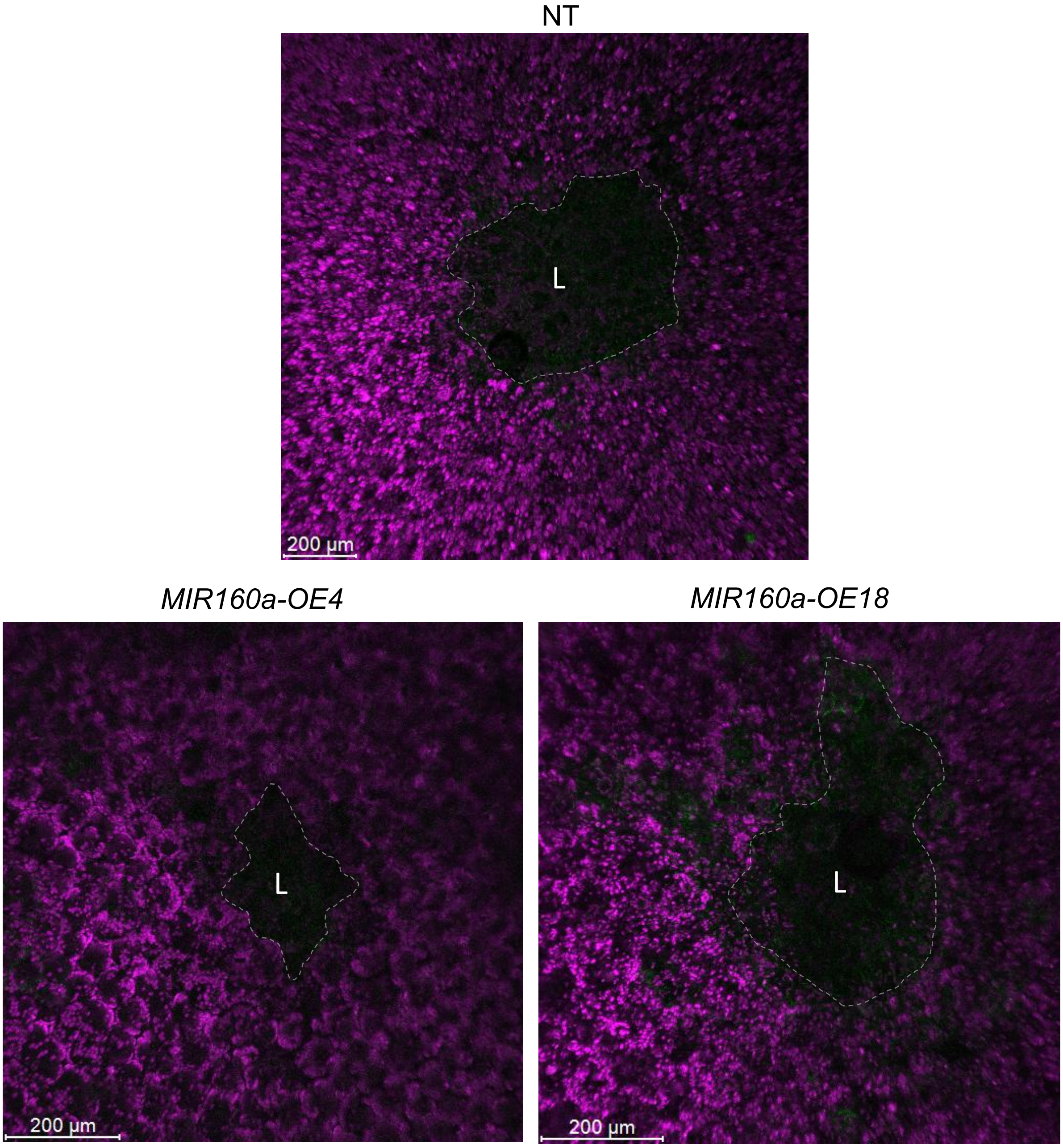


**Fig. S10: PVY remains confined to the cell death zone in miR160 overexpression plants.** PVY-N605(123)-GFP accumulation (green) was analyzed at fully developed lesion stage (at 6 days post inoculation) by confocal microscopy in miR160 overexpression lines (*MIR160a-OE4*, *MIR160a-OE18*) and in non-transgenic (NT) cv. Rywal. Figures represent the overlay of PVY-N605(123)-GFP accumulation (green) and chlorophyll fluorescence (purple). The lesion is marked by the white "L", and the edge of the cell death zone is marked by the light dotted line. Scale bar: 200 µm.

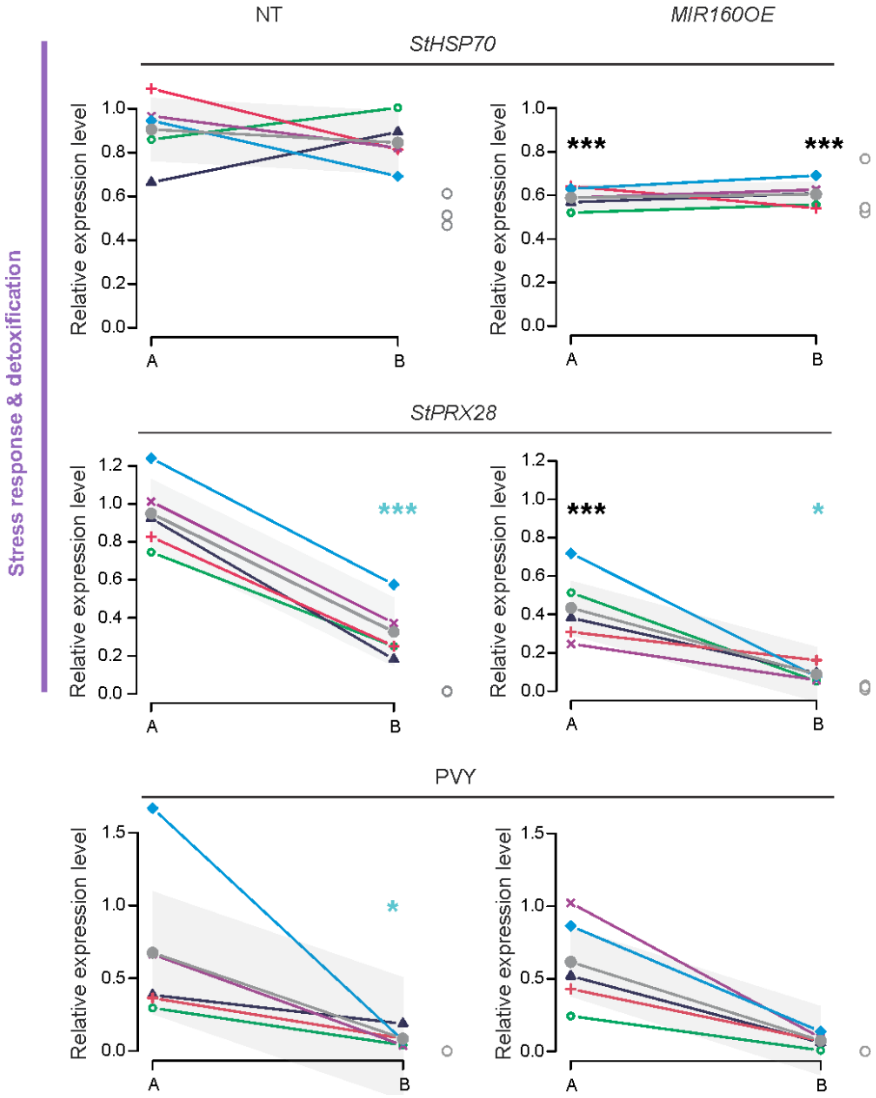


**Fig. S11: miR160 represses *StHSP70* and *StPRX28*.** Spatial response of *StHSP70*, *StPRX28* and PVY levels in lesion and its immediately adjacent cells (section A) and surrounding regions (section B). Expression values in mock (M)-inoculated tissue sections are shown as empty circles at the end of x-axis. Spatial profile models are shown as thick black lines, with 95% confidence interval bands presented in grey. Relative gene expression values of individual lesions are presented with colored symbols connected by a line. Black asterisks indicate statistically significant differences in gene expression in miR160 OE plants in sections A and B relative to NT (non-transgenic) cv. Rywal, whereas blue asterisks indicate statistically significant differences between sections A and B within each genotype. All significance levels were assessed by Tukey HSD test (*$P$<0.05, **$P$<0.01, *** $P$< 0.001).

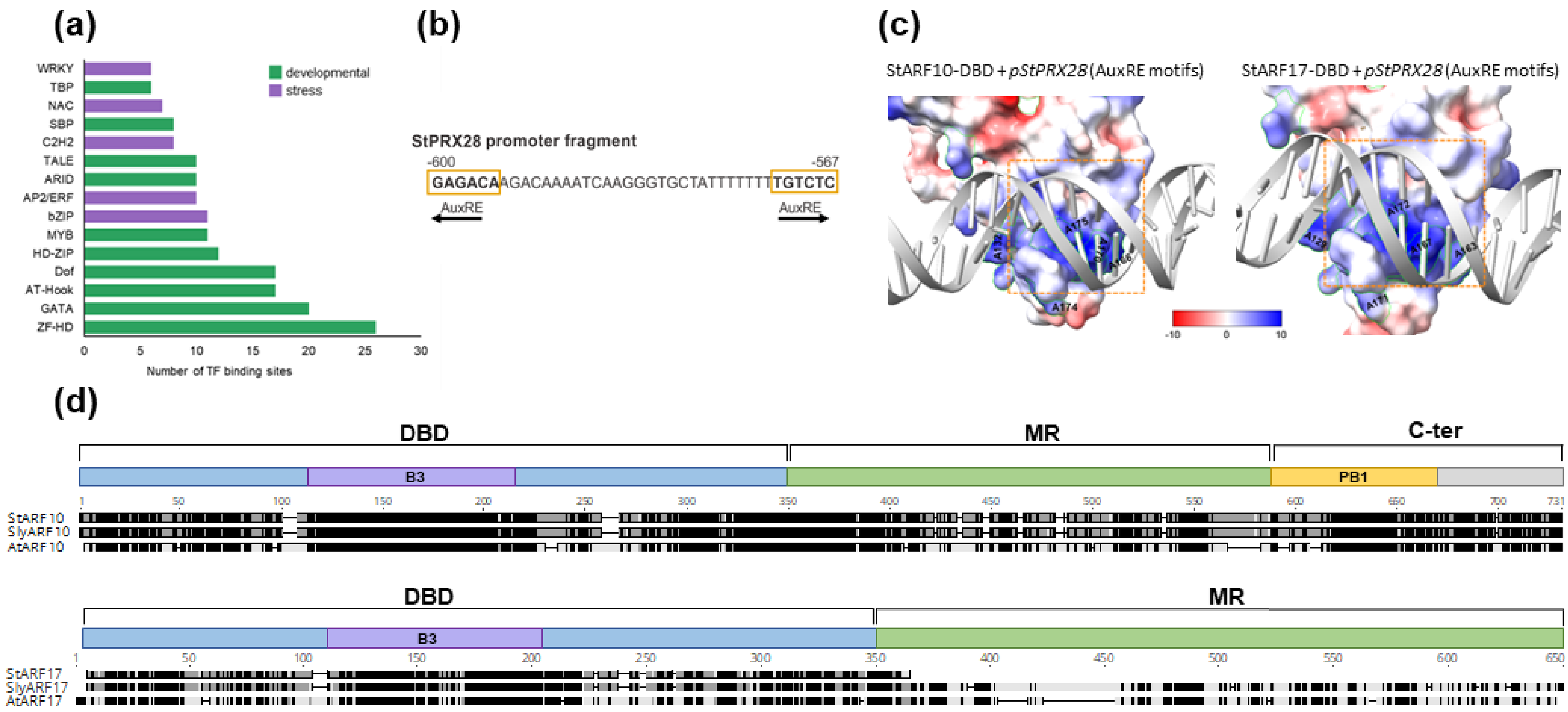


**Fig. S12**: **StARF10 and StARF17 recognize AuxRE in *StPRX28* promoter.** (a) Enriched transcription-factor (TF) binding motifs in the *StPRX28* promoter (*pStPRX28*). The promoter contains multiple motifs for TFs linked to development and salicylic acid/ethylene signaling. The top 15 motifs are shown and grouped by primary role in development or stress response. For the complete list of motifs see Table S12b. (b) *StPRX28* promoter contains two ARF-binding motifs (AuxRE) at 567 bp (+ strand) and at 600 bp (- strand) upstream of the transcription start site (yellow squares). (c) Structural models of StARF10-DBD and StARF17-DBD (DBD; DNA binding domains) binding to both identified AuxREs (orange squares). Both StARF10 and StARF17 displayed strongly positive ESP at the region binding to AuxRE within *StPRX28* promoter; Scale: blue-positive, red-negative ESP. (d) Protein sequence alignment of potato, tomato and Arabidopsis ARF10 and ARF17. A typical ARF protein possesses three regions: a conserved N-terminal DNA-binding domain (DBD; ~ 350 aa), followed by a nonconserved middle region (MR), and a C-terminal PB1 domain. The DBD domain contains a B3 DNA binding motif that binds specifically to AuxREs. Unlike ARF10, ARF17 lacks the C-terminal domain.

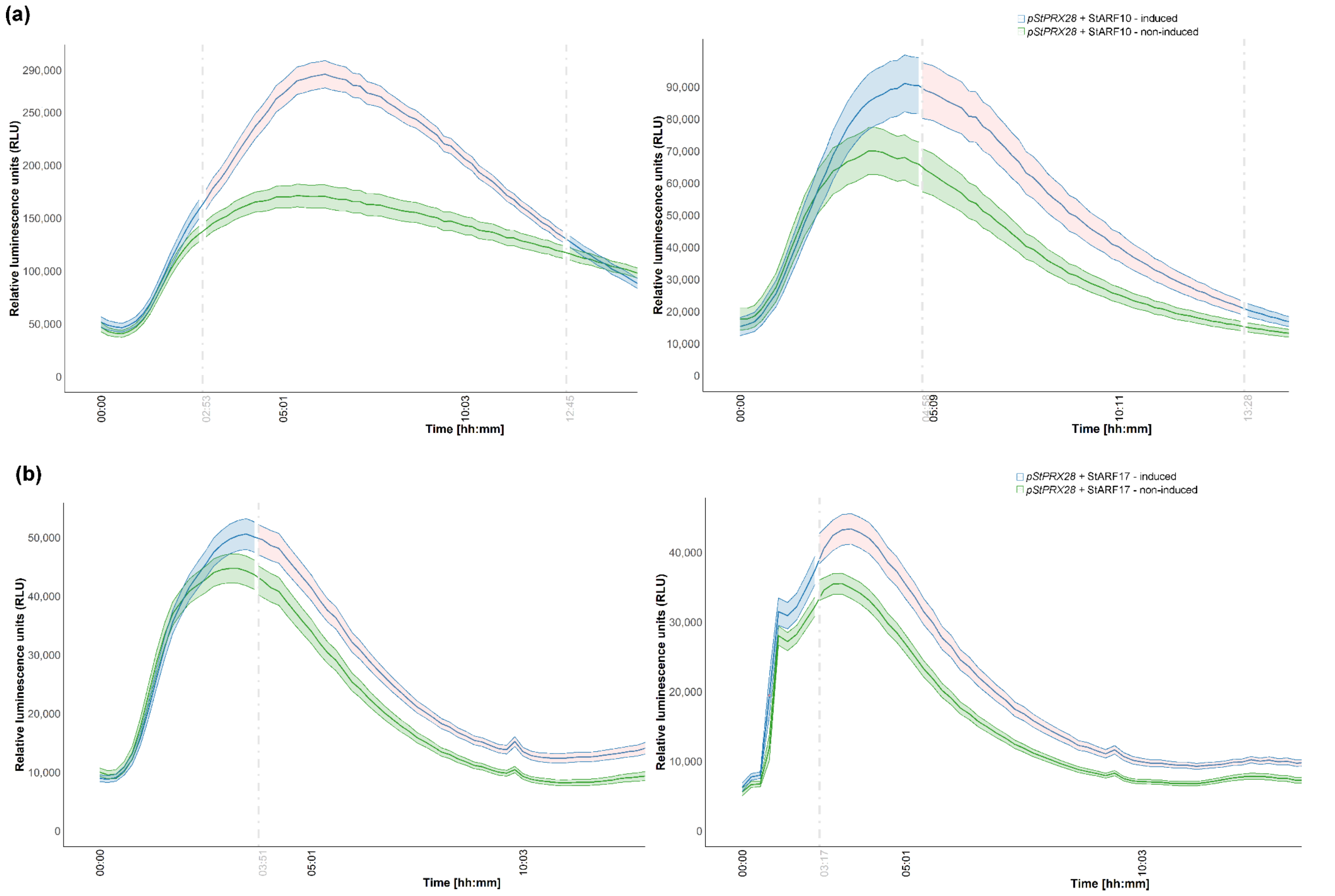
(a)
pStPRX28 + StARF10 - induced
pStPRX28 + StARF10 - non-induced
Relative luminescence units (RLU)
Time [hh:mm]
290,000
250,000
200,000
150,000
100,000
50,000
0
00:00
02:53
05:01
10:03
12:45
90,000
80,000
70,000
60,000
50,000
40,000
30,000
20,000
10,000
04:58
05:09
10:11
13:28
(b)
pStPRX28 + StARF17 - induced
pStPRX28 + StARF17 - non-induced
50,000
40,000
30,000
20,000
10,000
03:51
03:17

**Fig. S13: StARF10 and StARF17 activate expression of class III plant peroxidase *StPRX28*.** Transactivation assay results showing *in planta StPRX28* promoter (*pStPRX28*) activation by (a) StARF10 and (b) StARF17 based on two additional independent experiments for each construct. The β-estradiol inducible StARF10 and StARF17 effector vectors and the *StPRX28 promoter* fused to the firefly luciferase reporter were co-infiltrated in *N. benthamiana* leaves. Relative luminescence units (RLU) were measured over time in β-estradiol treated (blue) and untreated control (green) samples. Statistically significant differences between β-estradiol treated and untreated samples (pointwise group comparisons using pairwise Wilcoxon tests with Holm correction) are highlighted with change in ribbon color (light pink; $P < 0.05$). Solid thick lines within the graphs connect mean values for the treated/induced (blue) and untreated/non-induced samples (green). Solid thin lines indicate the ± standard error of the mean for the treated (blue) and untreated (green) samples.

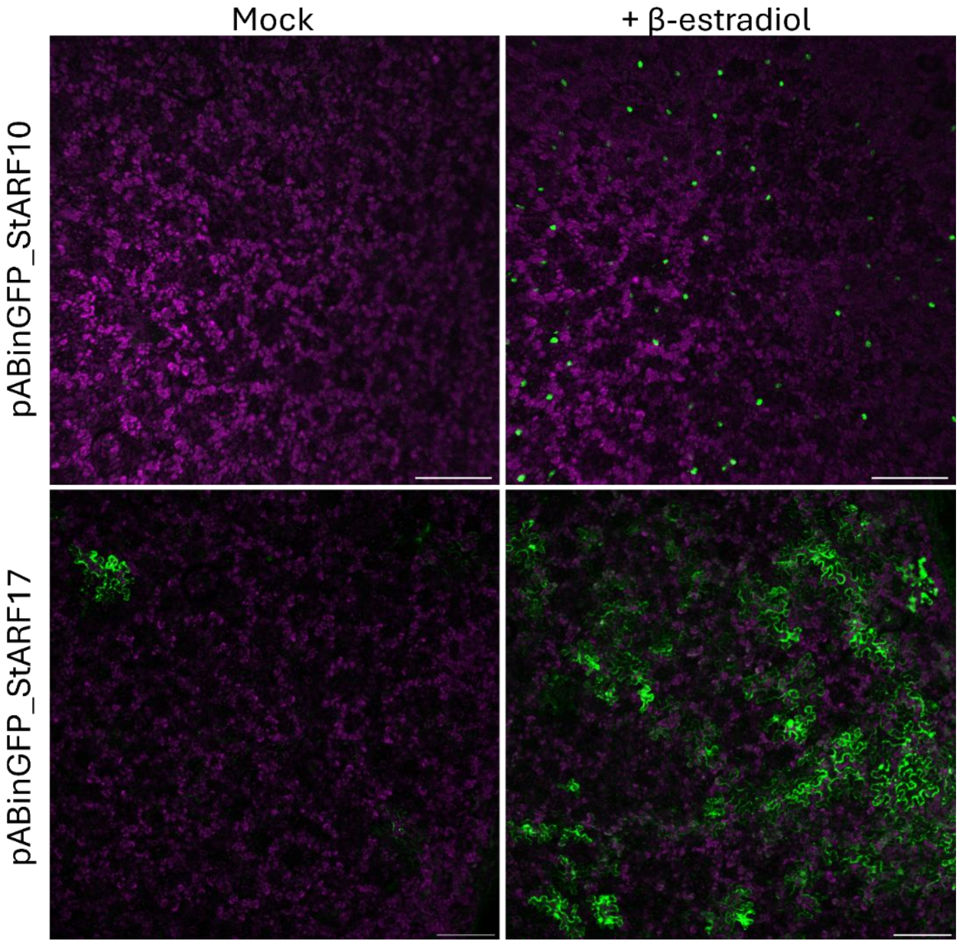


**Fig. S14: Confirmation of expression of GFP-tagged StARF10 and StARF17 under β-estradiol inducible promoter *in N. benthamiana* leaves.** Confocal microscopy images of the cells expressing GFP-tagged StARF10 (top) and GFP-tagged StARF17 (bottom) under the β-estradiol inducible system (pABinGFP). When the samples were treated with inducer (right images) the GFP signal is detected. Figures show strong signal of GFP-tagged StARF10 in the nuclei, whereas GFP-tagged StARF17 was detected in the nuclei and cytoplasm. Figures represent the overlay of GFP (green) and chlorophyll fluorescence (purple). Scale bar: 200 µm.

(a)

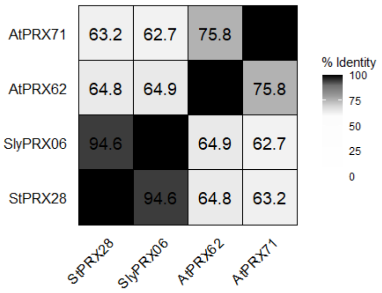


(b)

**Predicted localizations:** Extracellular
**Predicted signals:** Signal peptide

| Localization | Cytoplasm | Nucleus | Extracellular | Cell membrane | Mitochondrion | Plastid | Endoplasmic reticulum | Lysosome/Vacuole | Golgi apparatus | Peroxisome |
|---|---|---|---|---|---|---|---|---|---|---|
| Probability | 0.0863 | 0.0774 | 0.6953 | 0.1811 | 0.0164 | 0.1749 | 0.3786 | 0.5716 | 0.1128 | 0.0014 |

CELLO Prediction:

| | |
|---|---|
| Extracellular | 2.690 * |
| PlasmaMembrane | 1.220 |
| Mitochondrial | 0.273 |
| Cytoplasmic | 0.208 |
| Chloroplast | 0.196 |
| Vacuole | 0.147 |
| Nuclear | 0.096 |
| Peroxisomal | 0.057 |
| Lysosomal | 0.043 |
| ER | 0.037 |
| Golgi | 0.027 |
| Cytoskeletal | 0.005 |

(c)

**Prediction:** Signal Peptide (Sec/SPI)

Cleavage site between pos. 30 and 31. Probability 0.978424

| Protein type | Other | Signal Peptide (Sec/SPI) |
|---|---|---|
| Likelihood | 0.0002 | 0.9998 |

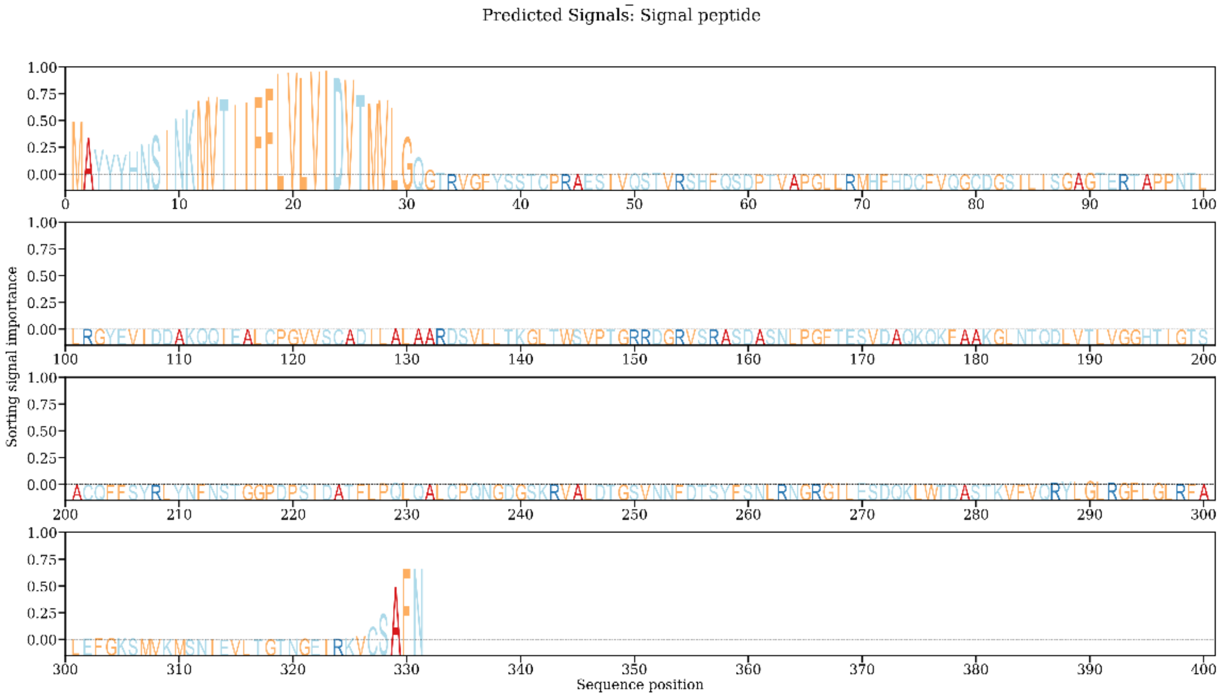

(d)

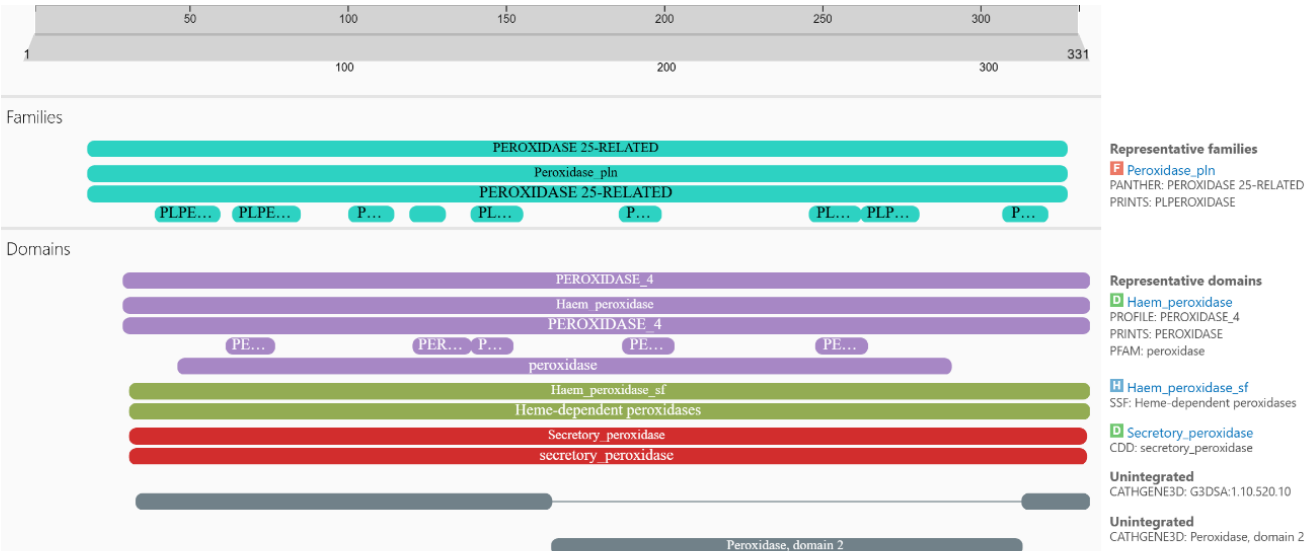


**Fig. S15: StPRX28 is Class III apoplastic peroxidase**. (a) Similarity color scheme of StPRX28 and its tomato and Arabidopsis orthologs, based on the identity score matrix computed using Clustal Omega web platform. (b) StPRX28 is predicted to be targeted to the extracellular space. Subcellular localization prediction using DeepLoc 2.0 and CELLO v2.5. (c) Signal peptide prediction of StPRX28 using SignalP v6.0 and DeepLoc 2.0. (d) StPRX28 contains a peroxidase domain, as well as heme- and calcium-binding sites. Domain predictions were performed using InterPro (https://www.ebi.ac.uk/interpro/).

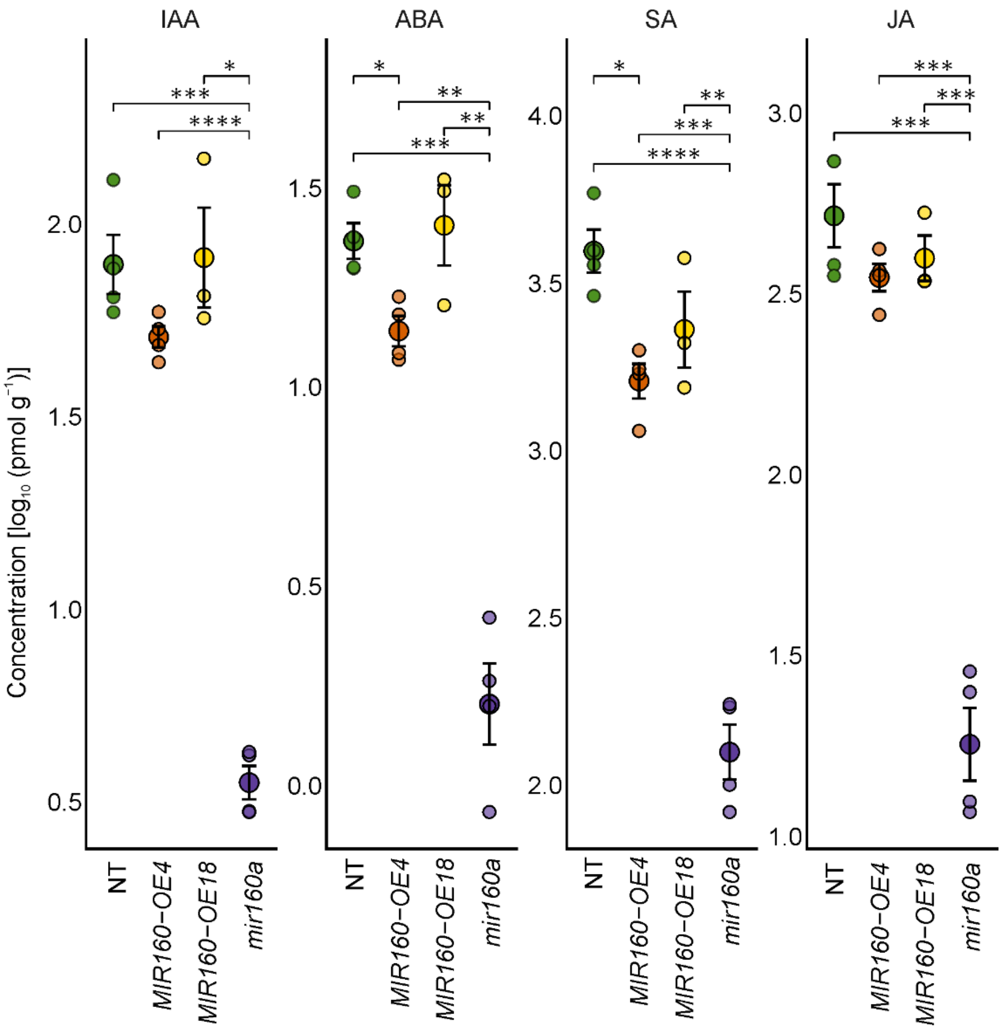


**Fig. S16: Hormone levels in fully developed lesions and surrounding tissue at 5 days post infection (dpi) in miR160-overexpression (*MIR160-OE4* and *MIR160-OE18*) and miR160 knockdown (*mir160a*) lines, relative to non-transgenic (NT) plants.** Arithmetic mean ± standard error of the mean is shown. Asterisks denote statistically significant differences based on Games Howell Post-hoc applied to $\log_{10}$-transformed values (*$P < 0.05$, **$P < 0.01$, ***$P < 0.001$). IAA: indole-3-acetic acid, ABA: abscisic acid, SA: salicylic acid, JA: jasmonic acid.

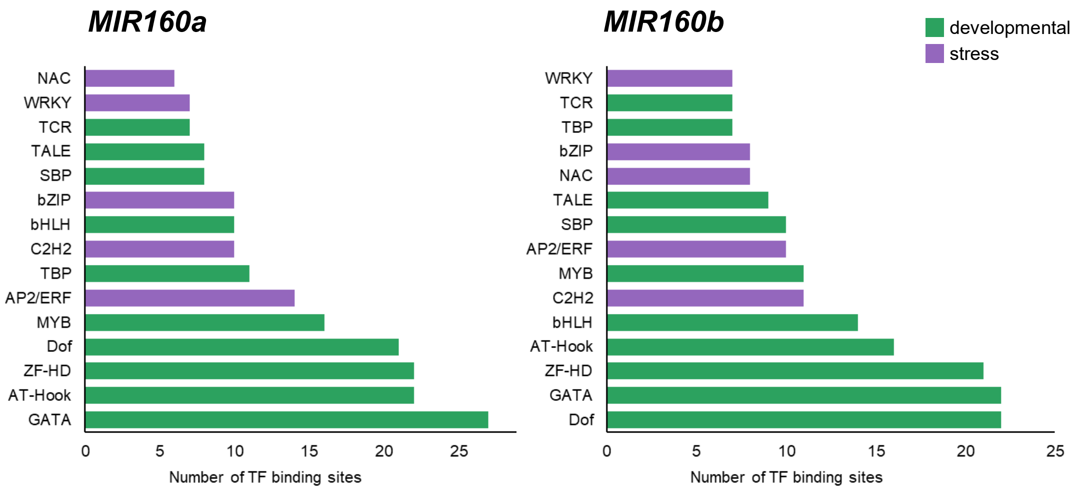


**Fig. S17: Enriched transcription-factor binding motifs in the *MIR160a* and *MIR160b* promoters**. The top 15 motifs are shown and grouped by primary role in development or stress response. For the complete list of motifs see Table S12b.

**Experiment No. 1**

**(a) Heat stress**

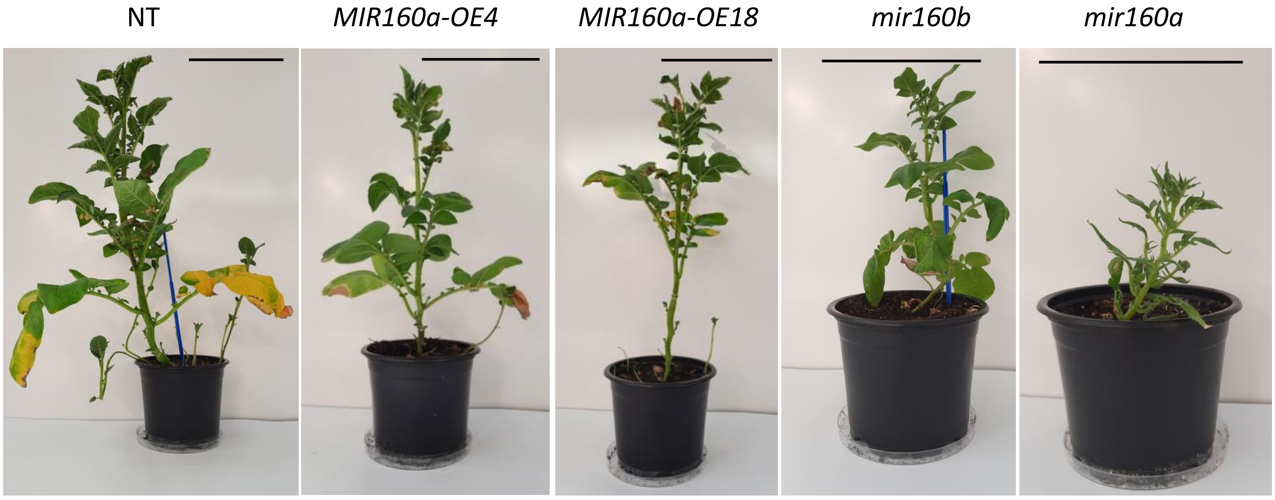


**(b) Control conditions**

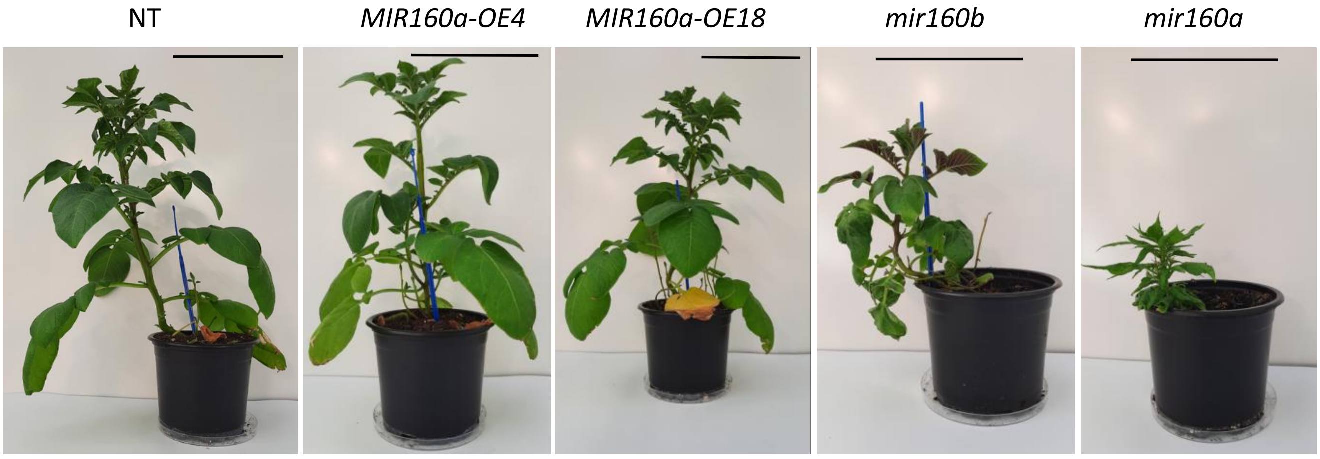


**Fig. S18: miR160 increases sensitivity to heat in potato (Experiment No. 1).** (a) Heat-stress phenotypes of miR160-overexpression lines (*MIR160a-OE-4* and *MIR160a-OE-18*), miR160 knockdown lines (*mir160b*, *mir160a*) and non-transgenic plants (NT) exposed to heat. NT and miR160 overexpression plants display heat induced necrosis and yellowing/senescence after two weeks of heat stress (30 °C during the day, 28 °C at night), whereas miR160 depleted plants show no symptoms. (b) The bottom panels show phenotypes of the same genotypes grown at 21 °C control conditions. Scale bar: 10 cm.

**Experiment No. 2**

**Heat stress, side view**

NT

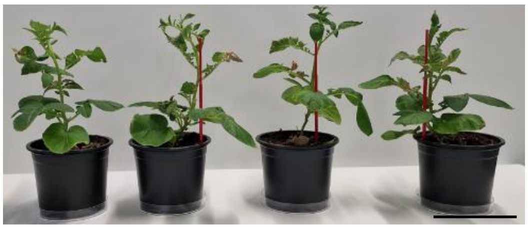

*MIR160a-OE4*

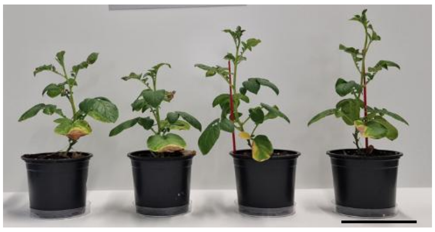

*mir160b*

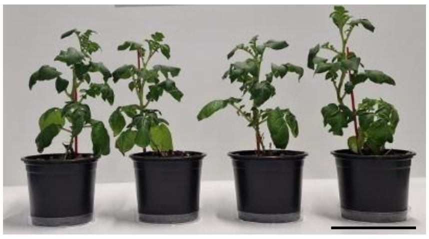

*MIR160a-OE18*

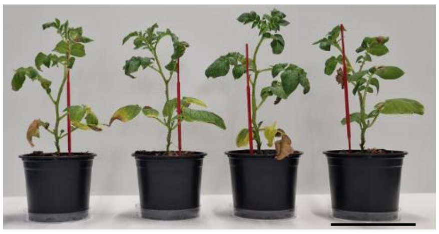

*mir160a*

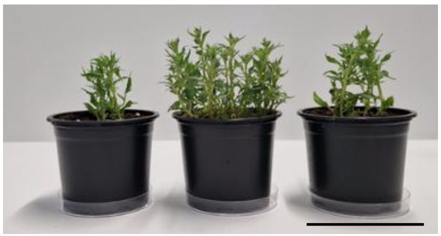

**Heat stress, top view**

NT

*MIR160a-OE4*

*MIR160a-OE18*

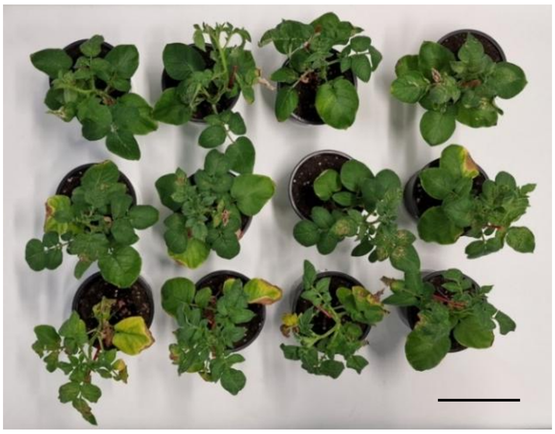

NT

*mir160b*

*mir160a*

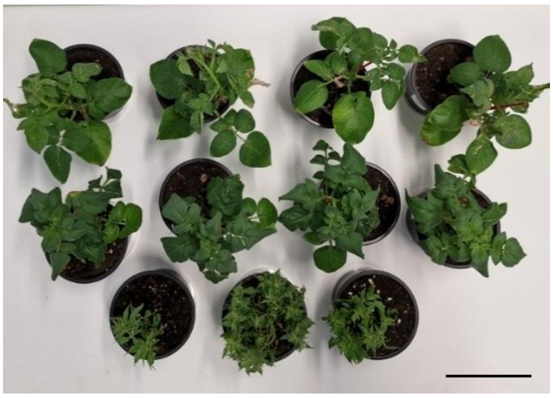

**Fig. S19: miR160 increases sensitivity to heat in potato (Experiment No. 2).** Heat-stress phenotypes of miR160-overexpression lines (*MIR160a-OE-4* and *MIR160a-OE-1*8), miR160 knockdown lines (*mir160b*, *mir160a*) and non-transgenic plants (NT) exposed to heat. NT and miR160 overexpression plants display heat induced necrosis and yellowing/senescence after two weeks of heat stress (30 °C during the day, 28 °C at night), whereas miR160 depleted plants show no symptoms. For control plants grown at 21 °C from the same experiment see Fig. S9. Scale bar: 10 cm.

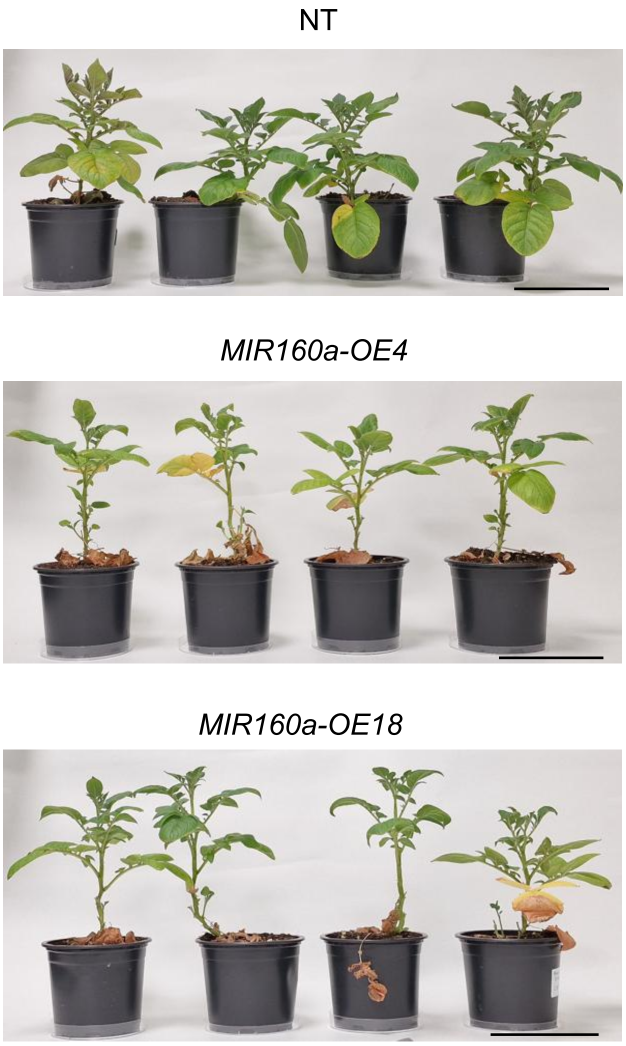


**Fig. S20: Increased miR160 levels are associated with accelerated senescence at later time points**. Phenotype of PVY-infected miR160 overexpression lines (*MIR160a-OE4*, *MIR160a-OE18*) and knockdown lines (*mir160a*, *mir160b*) four weeks post-inoculation, grown under normal conditions, compared with non-transgenic plants (NT) plants. Scale bar: 10 cm.

**(a)**

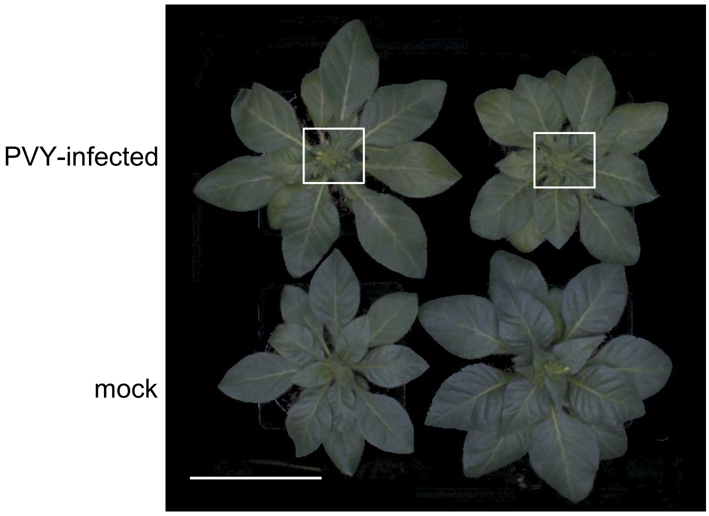


**(b)**

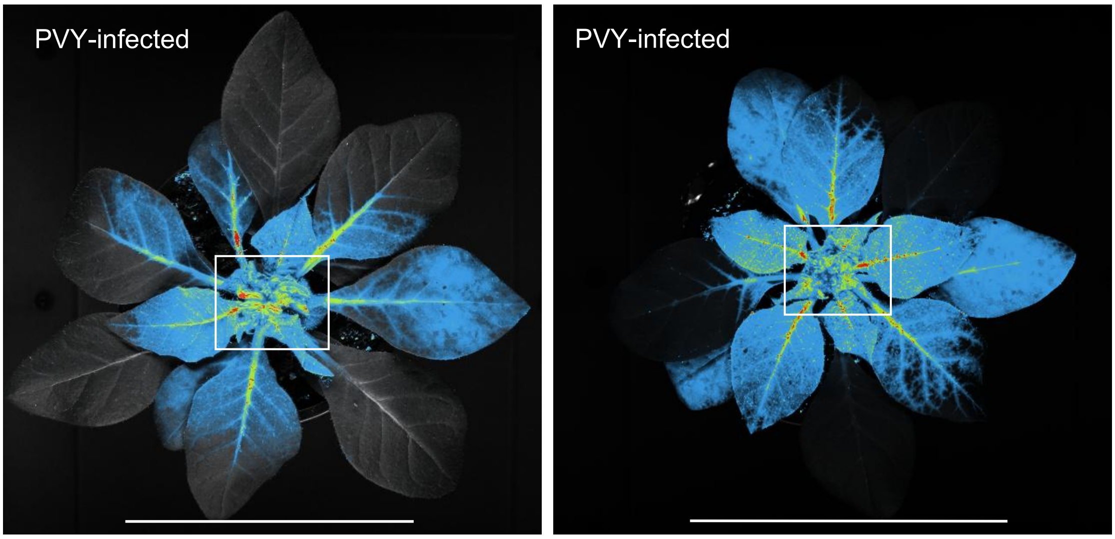


**Fig. S21: PVY induces leaf curling in *Nicotiana benthamiana.*** (a) Top view of plants inoculated with PVY-N605(123)-GFP or mock. The two plants in the upper row are PVY-infected, whereas the plants in the lower row are mock-inoculated controls. (b) Systemic distribution of PVY in infected plants, visualized by GFP fluorescence after two weeks post inoculation. The white squares indicate curled leaves. Scale bar: 10 cm.
